# A self-limiting orexin–habenula circuit for stress resilience

**DOI:** 10.64898/2026.05.07.722567

**Authors:** Soo Hyun Yang, Esther Yang, Jin Taek Jung, Jaekwang Lee, Gyeong Hee Pyeon, Yong Sang Jo, Hyung Sun Park, Ji Wook Moon, Jae–Yong Park, Kyung–Jun Boo, Dongmin Lee, Sungkun Chun, Hyeijung Yoo, Hyun Woo Lee, Hyun Kim

## Abstract

Resilience requires neural systems that mobilise active coping during stress while limiting its persistence to preserve homeostasis under sustained challenge^1–4^. Here we identify a self-limiting orexin–habenula circuit in which lateral hypothalamic orexin neurons engage aromatic L-amino acid decarboxylase-expressing D-neurons (encoded by *Ddc*) in the lateral habenula via orexin receptor type 2 (OX2R)^5–7^. Activation of this pathway increased nucleus accumbens dopamine and promoted active coping and positive valence. Optotagging revealed rapid stress-evoked recruitment followed by post-stress suppression. In lateral habenula neurons, orexin peptides exerted dissociable effects: orexin-A engaged an OX2R-dependent inhibitory programme superimposed on a parallel inward current, whereas orexin-B did not reproduce this inhibitory profile and instead exerted a distinct membrane effect. Chronic stress disrupted this buffering system through coordinated inflammatory activation, promoter methylation and erosion of D-neuron identity and orexin responsiveness. Restoring orexin-A reversed behavioural and molecular deficits through OX2R-dependent suppression of nuclear factor kappa B signalling, preservation of *Tet2* expression, and demethylation-linked maintenance of the *Ddc* programme. Together, these findings define a self-limiting orexin–habenula resilience circuit that enables adaptive coping while constraining stress-induced vulnerability.

## Main

Stress is a double-edged biological force. Acute challenges normally recruit adaptive processes that preserve behavioural flexibility and promote active coping, whereas chronic or uncontrollable stress increases vulnerability to psychiatric disorders, including depression^1–4^. This contrast implies the existence of neural mechanisms that not only mobilise coping responses during stress but also terminate them appropriately, thereby preventing adaptive responses from escalating into pathological states. However, the identity and operating principles of such self-limiting resilience circuits remain poorly understood.

The lateral habenula (LHb) is central to this question. Long regarded as an anti-reward hub, the LHb is classically thought to suppress midbrain dopaminergic function and promote aversion, helplessness, and depression-related behavioural states^8–14^. However, this framework leaves an important paradox unresolved: acute stress frequently activates the LHb^15^ yet does not ordinarily produce depression^5,16,17^.

In our previous work, we identified a molecularly distinct LHb subpopulation—D-neurons expressing aromatic L-amino acid decarboxylase (AADC, also known as dopa decarboxylase; encoded by *Ddc* in mouse and *DDC* in human)—and showed that, during acute stress, these neurons engage a protective trace-aminergic mechanism that suppresses rostromedial tegmental nucleus (RMTg) GABAergic neurons, sustains ventral tegmental area (VTA) dopaminergic tone, and supports active coping^5–7,13,18,19^. These findings suggested that the LHb is not uniformly depressogenic but instead contains a resilience-associated module capable of counterbalancing its canonical anti-reward output.

Epigenetic mechanisms are increasingly recognised as key regulators of stress susceptibility and resilience, linking inflammatory signalling to stable transcriptional reprogramming in the brain. In particular, ten-eleven translocation 2 (Tet2) has been implicated in activity-dependent DNA demethylation, inflammatory gene regulation, and the maintenance of neuronal transcriptional identity^20^, raising the possibility that disruption of Tet2-dependent pathways may contribute to the erosion of stress-adaptive neural programmes.

These observations prompted two central questions. First, what upstream input recruits LHb D-neurons during stress, and how is their coping-promoting activity subsequently constrained? Second, why does this protective programme fail under chronic stress, and by what mechanism is *Ddc* expression reduced in depression-like states? Resolving these questions is essential for understanding not only how adaptive stress responses are organised in the healthy brain but also how the same circuit architecture becomes destabilised under prolonged challenge.

The lateral hypothalamic area (LHA) is ideally situated to orchestrate this process. Orexin is a particularly compelling candidate signal because of its established roles in arousal, motivated behaviour and stress adaptation as well as its broader influence on inflammatory and homeostatic state regulation^21–25^. We therefore investigated whether LHA orexin inputs engage and stabilise an LHb D-neuron resilience module.

We identify an orexin LHA→LHb pathway that recruits D-neurons enriched in orexin receptor type 2 (OX2R; encoded by *Hcrtr2* in mouse and *HCRTR2* in humans), promotes active coping and exhibits biphasic dynamics consistent with a self-limiting stress–relief architecture. We further show that chronic stress disrupts this module through inflammatory and epigenetic erosion of D-neuron identity, whereas restoring orexin-A rescues D-neuron function through OX2R-dependent inflammatory and epigenetic stabilisation.

### LHA orexin inputs to LHb D-neurons

To identify upstream circuitry that engages LHb D-neurons, we mapped monosynaptic afferent inputs using retrograde tracing with Cre-dependent helper AAVs (TVA and RG) and EnvA-pseudotyped glycoprotein deleted rabies virus (RVΔG) in *Ddc*^Cre^ mice (Fig. 1a and Extended Data Fig. 1a). Brain-wide mapping revealed distributed presynaptic populations across multiple forebrain and hypothalamic regions (Fig. 1b and Extended Data Fig. 1b,c). Guided by prior work implicating several stress-responsive LHb-projecting nodes^26,27^—including the medial septum, lateral septum, lateral preoptic area, LHA and entopeduncular nucleus—we focused subsequent analyses on these five regions (Fig. 1c). Among these, the LHA contained the most prominent RVΔG-labelled presynaptic population (Fig. 1c). Within the LHA, RVΔG-labelled presynaptic neurons overlapped with orexin immunoreactivity, indicating that a subset of hypothalamic inputs to LHb D-neurons arises from orexin neurons (Fig. 1d). Consistently, fluorescence *in situ* hybridisation (FISH) analysis showed that RVΔG-labelled LHA input neurons co-expressed *Hcrt* and *Vglut2*, indicating that these afferents include orexinergic glutamatergic neurons (Extended Data Fig. 1d–j).

**Figure 1.**
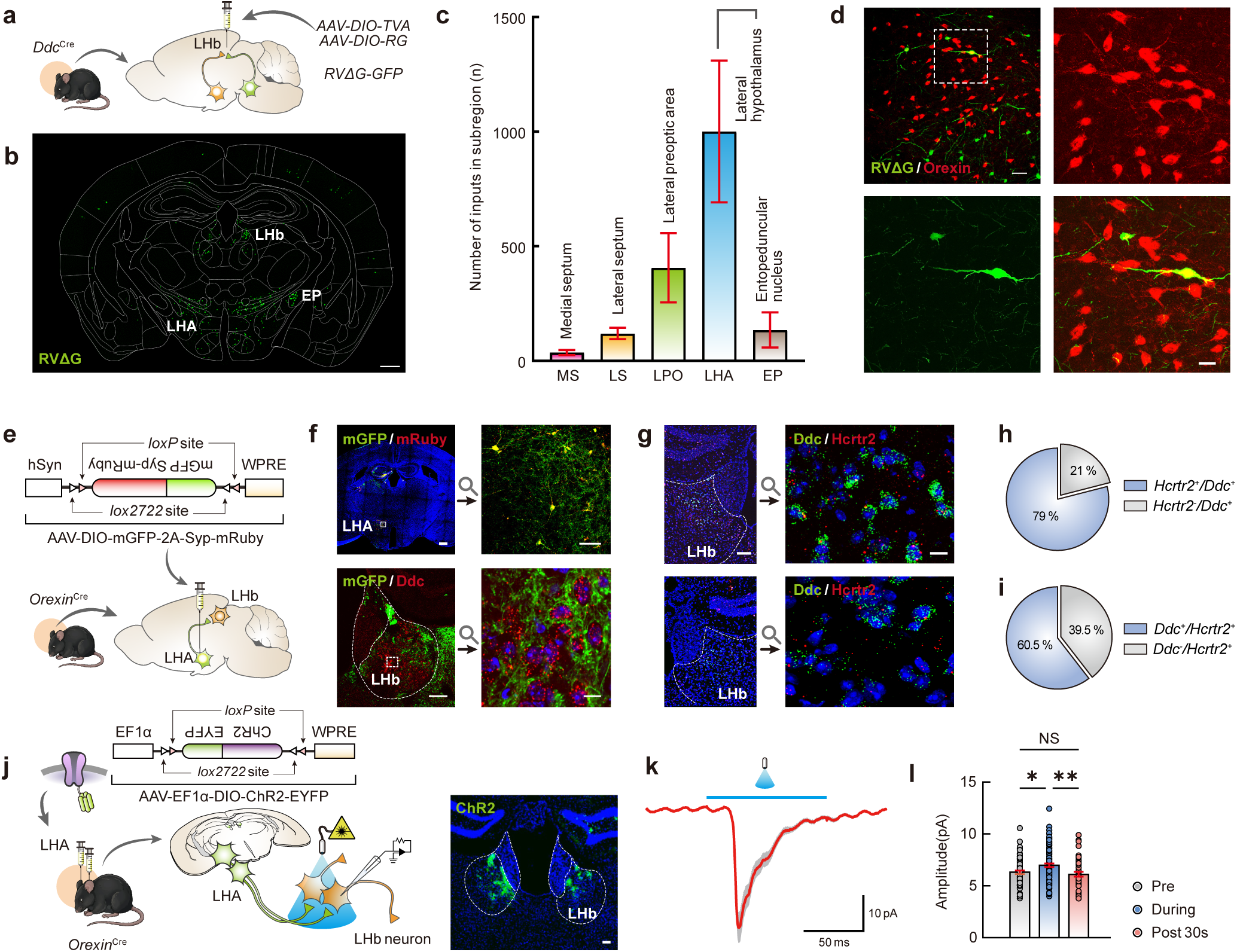
LHA orexin inputs define and functionally engage LHb D-neurons. **a**–**c**, Experimental schematic for monosynaptic retrograde tracing from LHb D-neurons in *Ddc*^Cre^ mice using Cre-dependent helper AAVs and RVΔG (**a**), brain-wide distribution of presynaptic inputs to LHb D-neurons (**b**), and quantification of candidate upstream regions projecting to LHb D-neurons (**c**). **d**, Representative images showing overlap between RVΔG-labelled presynaptic neurons and orexin-immunoreactive neurons in the LHA. **e**,**f**, Intersectional labelling strategy for LHA orexin projections in *Orexin*^Cre^ mice (**e**), and representative images showing dense orexin-positive fibres and presynaptic puncta in the LHb (**f**). **g**–**i**, Representative FISH images showing co-expression of *Ddc* and *Hcrtr2* in LHb neurons (**g**), quantification of the proportions of *Hcrtr2*^+^/*Ddc*^+^ and *Hcrtr2*⁻/*Ddc*^+^ cells among *Ddc*^+^ neurons (**h**), and quantification of the proportions of *Ddc*^+^/*Hcrtr2*^+^ and *Ddc*⁻/*Hcrtr2*^+^ cells among *Hcrtr2*^+^ neurons (**i**). **j**–**l**, Experimental schematic and representative viral expression image for optogenetic stimulation of LHA orexin terminals in acute LHb slices (**j**), representative light-evoked inward current trace (**k**), and quantification of current amplitude before, during and after photostimulation in LHb neurons (**l**). AAV, adeno-associated virus; RVΔG, glycoprotein-deleted rabies virus; LHA, lateral hypothalamic area; LHb, lateral habenula; FISH, fluorescence *in situ* hybridisation; *Ddc*, dopa decarboxylase; *Hcrtr2*, hypocretin receptor 2; EPSC, excitatory postsynaptic current. Error bars indicate mean ± s.e.m. NS, not significant; \**P* < 0.05, \*\**P* < 0.01. Scale bars, **b**: 500 μm; **d**: 50 μm (top) and 20 μm (bottom); **f**: 500 μm (top, left), 100 μm (top, right), 100 μm (bottom, left), 10 μm (bottom, right); **g**: 100 μm (left), 10 μm (right); **j**: 100 μm. Exact *n* and *P* values are provided in Supplementary Table 5.

We next validated direct orexin projections to the LHb using an intersectional viral strategy in *Orexin*^Cre^ mice, which revealed dense orexin^+^ fibres and presynaptic puncta within the LHb (Fig. 1e,f). FISH analysis further showed that most *Ddc*^+^ neurons co-expressed *Hcrtr2* (79.0%; Fig. 1g,h), whereas a substantial fraction of *Hcrtr2*^+^ neurons was also *Ddc^+^* (60.5%; Fig. 1g,i). Together, these data define an anatomical and receptor-enriched substrate whereby LHA orexin inputs recruit LHb D-neurons.

To test whether this projection is functionally coupled to LHb neurons, we expressed channelrhodopsin 2 (ChR2) in LHA orexin neurons in *Orexin*^Cre^ mice and optogenetically stimulated orexin terminals in acute LHb slices while performing whole-cell patch-clamp recordings from identified LHb neurons. Photostimulation evoked immediate inward currents, supporting direct functional connectivity between LHA orexin terminals and LHb D-neurons (Fig. 1k,l and Extended Data Fig. 1k,l).

### LHA–LHb orexin activation promotes coping

We next asked whether engaging LHA orexin input to OX2R-expressing LHb D-neurons is sufficient to promote active coping. In *Orexin*^Cre^ mice, projection-defined chemogenetic activation of LHb-projecting orexin neurons expressing hM3Dq (Fig. 2a) increased *Fos* expression in LHA orexin neurons and LHb D-neurons, reduced *Fos* expression in RMTg GABAergic neurons, and increased *Fos* expression across the mesolimbic pathway, including VTA dopaminergic neurons and nucleus accumbens (NAc) GABAergic neurons (Fig. 2b–f). Consistent with disinhibition of mesolimbic output, fibre photometry revealed a sustained (>1 h) elevation of NAc dopamine following intraperitoneal administration of clozapine N-oxide (CNO; Fig. 2g,h and Extended Data Fig. 2a,b). Behaviourally, the same manipulation increased active coping in the tail suspension test (TST; Fig. 2i–l) and forced swim test (FST; Fig. 2m–p) and induced conditioned place preference (CPP; Fig. 2q–s), indicating positive motivational valence.

**Figure 2.**
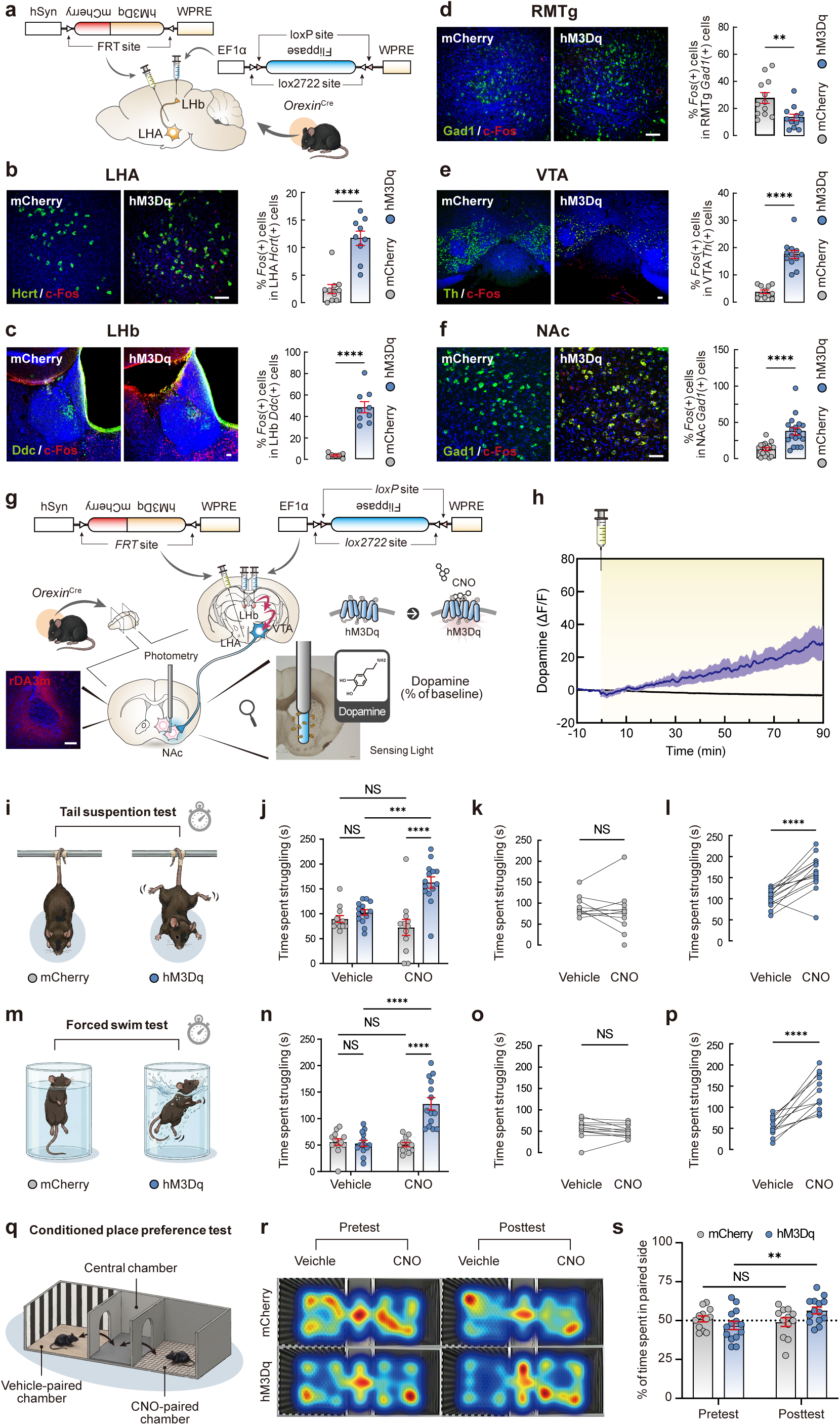
Activation of the LHA→LHb orexin pathway promotes coping and mesolimbic recruitment. **a**, Experimental schematic for projection-defined chemogenetic activation of LHb-projecting orexin neurons in *Orexin*^Cre^ mice. **b**–**f**, Representative *Fos* images and quantification showing increased activity in LHA orexin neurons (**b**), LHb D-neurons (**c**), reduced activity in RMTg GABAergic neurons (**d**), and increased activity in VTA dopaminergic neurons (**e**) and NAc GABAergic neurons (**f**) following chemogenetic activation. **g**,**h**, Fibre photometry schematic for monitoring NAc dopamine signalling following CNO administration (**g**), and representative trace with quantification showing sustained elevation of NAc dopamine (**h**). **i**–**l**, TST schematic (**i**), time spent struggling across groups (**j**), and paired comparisons of TST performance before and after CNO in mCherry control (**k**) and hM3Dq (**l**) mice. **m**–**p**, FST schematic (**m**), time spent struggling across groups (**n**), and paired comparisons of FST performance before and after CNO in mCherry control (**o**) and hM3Dq (**p**) mice. **q**–**s**, Experimental schematic for the CPP assay (**q**), representative heat maps from pre-test and post-test sessions (**r**), and quantification of time spent in the CNO-paired side (**s**). LHA, lateral hypothalamic area; LHb, lateral habenula; RMTg, rostromedial tegmental nucleus; VTA, ventral tegmental area; NAc, nucleus accumbens; CNO, clozapine N-oxide; TST, tail suspension test; FST, forced swim test; CPP, conditioned place preference. Error bars indicate mean ± s.e.m. NS, not significant; \*\**P* < 0.01, \*\*\**P* < 0.001, \*\*\*\**P* < 0.0001. Scale bars, **b**–**f**: 50 μm, **g**: 300 μm. Exact *n* and *P* values are provided in Supplementary Table 5.

To test sufficiency with greater temporal precision, we photostimulated orexin terminals in the LHb in *Orexin*^Cre^::ChR2 mice. Photostimulation rapidly increased struggling in a within-session OFF–ON–OFF design and induced real-time place preference (RTPP; Extended Data Fig. 3a,b). During tail suspension, mice began moving immediately when the light was turned on and stopped moving when it was turned off (Extended Data Fig. 3c,d). In the RTPP assay, ChR2 mice showed a significant preference for the photostimulation-paired chamber, whereas control mice did not (Extended Data Fig. 3e,f). These rapid behavioural transitions indicate that orexin-driven coping responses are tightly gated and rapidly reversible, consistent with the existence of an active counter-regulatory process (Extended Data Fig. 3d). Neither chemogenetic activation nor LHb terminal photostimulation altered locomotor measures, as assessed by total distance moved and mean velocity in the open field test (OFT) and RTPP assays (Extended Data Fig. 2c–h and Extended Data Fig. 3g–j), indicating that the place-preference effects were not attributable to a general increase in locomotor activity. Together, these results show that activating orexin input to the LHb is sufficient to elevate NAc dopamine and bias behaviour towards active coping under stress.

Because LHA orexin neurons are glutamatergic, these rapid behavioural effects are likely to reflect combined actions of fast glutamatergic transmission and orexinergic modulation; the D-neuron-specific OX2R knockdown and rescue experiments presented below define a specific contribution of orexin signalling within this broader projection system.

### Differential LHb actions of orexin

Because preprohypocretin (prepro-orexin) is processed into two endogenous peptides, orexin-A and orexin-B^28,29^, we next asked whether these peptides exert distinct synaptic and membrane actions in LHb neurons. Given that OX2R can couple to both pertussis toxin (PTX)-sensitive Gαi/o and PTX-insensitive Gαq pathways^30–32^, we examined the respective effects of orexin-A and orexin-B in acute LHb slices during whole-cell recordings (Fig. 3a). In voltage-clamp recordings performed in the presence of bicuculline and picrotoxin, such that GABAergic transmission was blocked throughout baseline and drug application, orexin-A did not significantly alter the excitatory postsynaptic current (EPSC) frequency (Fig. 3b,c) but significantly reduced the EPSC amplitude (Fig. 3d,e). This reduction in EPSC amplitude was prevented by the OX2R antagonist TCS-OX2-29 (Fig. 3d,e), indicating that orexin-A suppresses the excitatory synaptic strength in LHb neurons in an OX2R-dependent manner. In separate voltage-clamp recordings, orexin-A also significantly increased inward current relative to the baseline (Fig. 3f,g), revealing a parallel direct membrane action.

**Figure 3.**
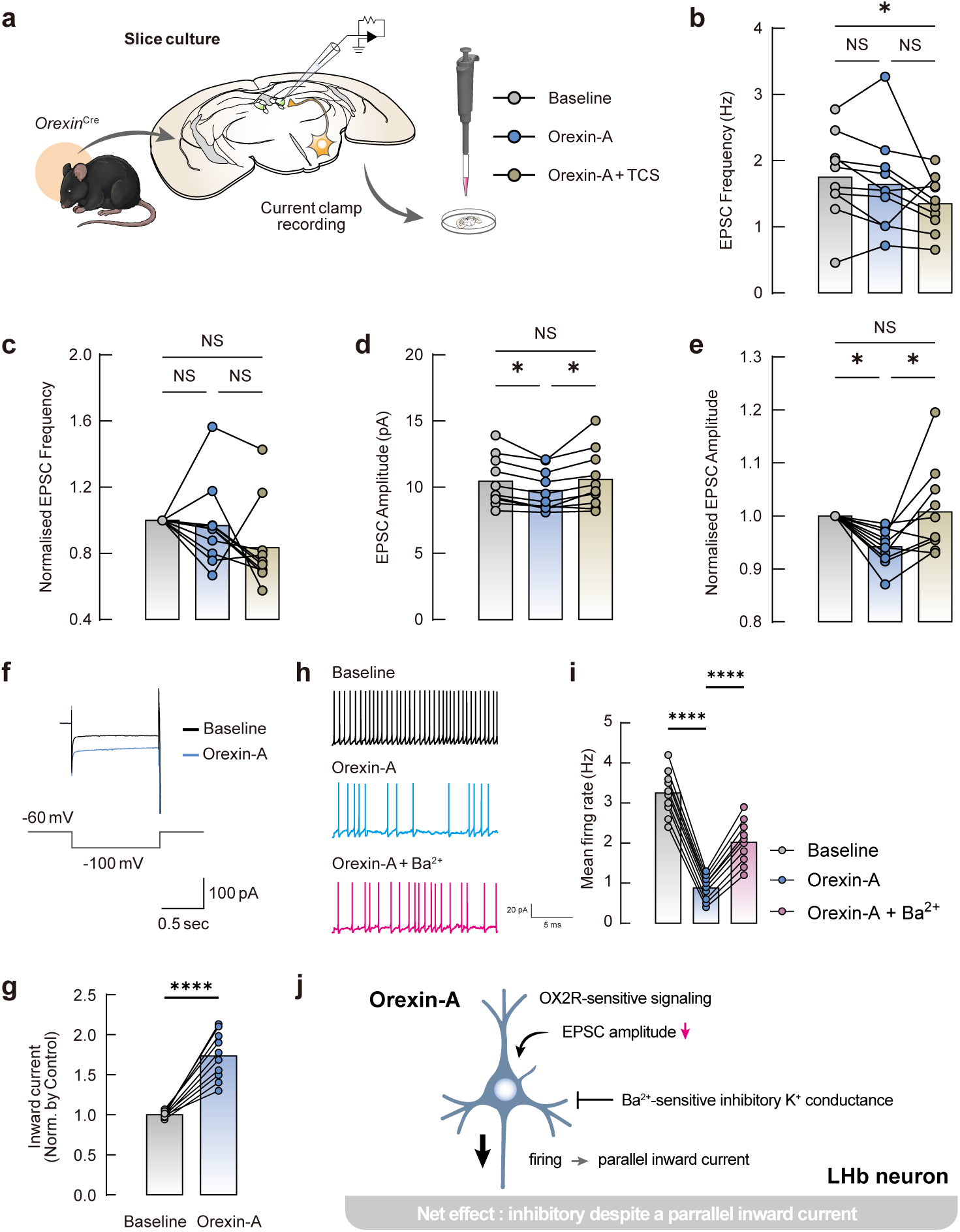
Orexin-A engages an OX2R-dependent inhibitory programme in LHb neurons. **a**, Experimental schematic for acute slice pharmacology and whole-cell recordings in LHb neurons. **b**,**c**, EPSC frequency (**b**) and normalised EPSC frequency (**c**) in the presence of bicuculline and picrotoxin after orexin-A application. **d**,**e**, EPSC amplitude (**d**) and normalised EPSC amplitude (**e**), showing orexin-A-induced suppression of excitatory synaptic strength and reversal by the OX2R antagonist TCS-OX2-29. **f**,**g**, Representative voltage-clamp trace showing orexin-A-induced inward current (**f**) and quantification of the inward current amplitude (**g**). **h**,**i**, Representative current-clamp traces showing that orexin-A reduces firing relative to baseline but increases firing in the presence of Ba^2+^ (**h**), and quantification of firing rate across baseline, orexin-A and orexin-A+Ba^2+^ conditions (**i**). **j**, Working model summarising orexin-A action in LHb neurons, in which OX2R-mediated signalling reduces EPSC amplitude and recruits a Ba^2+^-sensitive inhibitory K^+^ conductance that overcomes a parallel inward current, resulting in a net inhibitory effect on neuronal firing. LHb, lateral habenula; EPSC, excitatory postsynaptic current; OX2R, orexin receptor 2; K^+^, potassium. Error bars indicate mean ± s.e.m. NS, not significant; \**P* < 0.05, \*\*\*\**P* < 0.0001. Exact *n* and *P* values are provided in Supplementary Table 5.

To determine how these effects converge on neuronal output, we performed current-clamp recordings in normal artificial cerebrospinal fluid. Under these conditions, orexin-A significantly reduced firing relative to baseline (Fig. 3h,i). However, Ba^2+^ converted the orexin-A response from firing suppression to increased firing relative to baseline (Fig. 3h,i), indicating that the net inhibitory effect of orexin-A depends on a Ba^2+^-sensitive K^+^ conductance downstream of OX2R signalling.

In contrast, orexin-B did not significantly alter the EPSC frequency or amplitude under the same GABA-blocked conditions, and no TCS-OX2-29-sensitive effect was observed. Thus, unlike orexin-A, orexin-B did not measurably suppress excitatory synaptic strength in LHb neurons (Extended Data Fig. 4a–d). In separate voltage-clamp recordings, orexin-B reduced the inward-current response relative to baseline (Extended Data Fig. 4e,f), indicating a membrane effect distinct from that of orexin-A. Nevertheless, in current-clamp recordings, orexin-B increased spike output across injected current steps (Extended Data Fig. 4g,h), and this increase was further enhanced in the presence of Ba^2+^ (Extended Data Fig. 4i,j). These findings indicate that orexin-B does not reproduce the OX2R-dependent, Ba^2+^-sensitive inhibitory profile observed with orexin-A and instead exerts a distinct net effect on LHb neuronal output, the underlying membrane mechanism of which remains unresolved.

Together, these results identify dissociable peptide-dependent actions of orexin signalling in the LHb: orexin-A engages an OX2R-dependent reduction in EPSC amplitude together with a Ba^2+^-sensitive inhibitory influence on firing (Fig. 3j), whereas orexin-B does not reproduce this inhibitory profile and instead distinctly alters membrane properties while increasing neuronal firing.

### Biphasic orexin dynamics under stress

Having found that orexin-A exerts a net inhibitory influence on LHb neuronal output, we next asked how LHb-projecting orexin neurons dynamically regulate their output during stress *in vivo*. To address this, we performed projection-defined optotagging/optrode recordings from LHb-projecting orexin neurons in the LHA of *Orexin*^Cre^ mice, identifying units based on reliable, time-locked light-evoked spiking (10 pulses at 30 Hz; Fig. 4a–c and Extended Data Fig. 5a).

**Figure 4.**
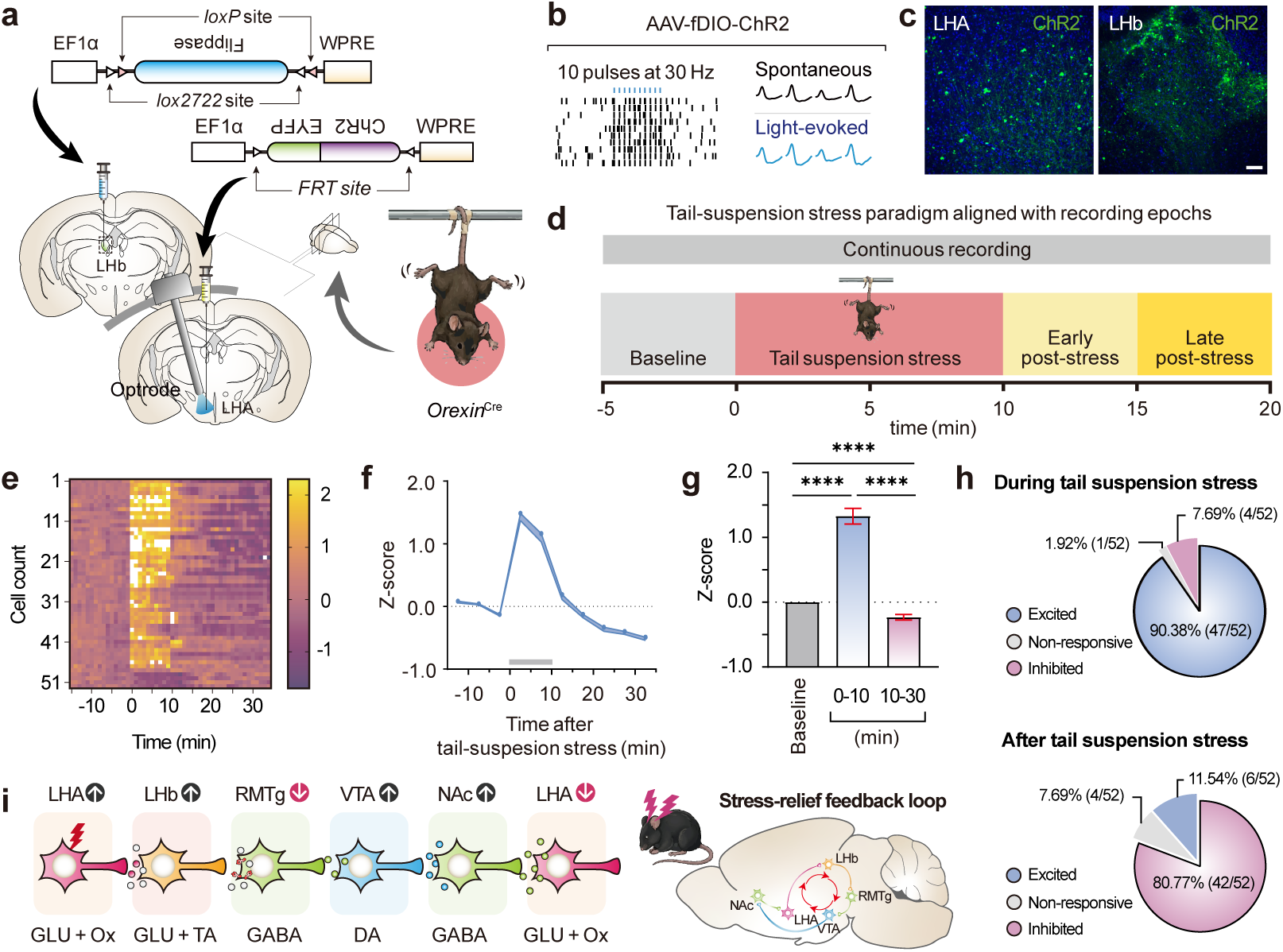
LHb-projecting orexin neurons exhibit biphasic dynamics during stress. **a**–**d**, Experimental schematic for projection-defined optotagging and in vivo optrode recording of LHb-projecting orexin neurons in *Orexin*^Cre^ mice (**a**), optical identification of ChR2-expressing LHb-projecting orexin neurons by light-evoked spiking (**b**), representative images of viral expression in the LHA and LHb (**c**), and tail-suspension stress paradigm aligned with recording epochs (**d**). **e**–**g**, Heat map of normalised firing in optotagged LHb-projecting orexin neurons aligned to tail-suspension stress (**e**), population-average normalised firing across recording epochs (**f**), and comparisons of average firing rates before stress (baseline), during tail suspension (0–10 min), and after tail suspension (10–30 min) (**g**). **h**, Proportions of optotagged LHb-projecting orexin neurons showing excitation, no response, or inhibition during and after tail-suspension stress. **i**, Working model of a stress–relief feedback loop: LHA→LHb orexin neurons transiently engage an LHb–RMTg–VTA–NAc coping pathway during stress, which is subsequently curtailed through feedback within the broader LHA–LHb–RMTg–VTA–NAc–LHA network. LHA, lateral hypothalamic area; LHb, lateral habenula; RMTg, rostromedial tegmental nucleus; VTA, ventral tegmental area; NAc, nucleus accumbens; ChR2, channelrhodopsin-2; GLU, glutamate; Ox, orexin; TA, trace amine; DA, dopamine. Error bars indicate mean ± s.e.m. NS, not significant; \*\*\*\**P* < 0.0001. Scale bars, **c**: 50 μm. Exact *n* and *P* values are provided in Supplementary Table 5.

During a 10-min tail-suspension stress, LHb-projecting orexin neurons displayed a biphasic activity profile. Across the recorded population, spiking increased rapidly at stress onset and peaked during the early phase of the suspension period (Fig. 4e–g and Extended Data Fig. 5b,c). As the stress episode progressed—and particularly after suspension ended—activity declined, with many units showing suppression below baseline rather than returning to baseline (Fig. 4g). Consistent with this pattern, most optotagged LHb-projecting orexin neurons exhibited stress-evoked activation (47/52 units), and the majority also showed a subsequent suppressive phase (42/52 units; Fig. 4h).

Together with the *Fos*-defined downstream recruitment pattern and prior evidence for NAc-to-LHA feedback^33–35^, these dynamics are consistent with a stress–relief feedback model in which LHA→LHb orexin neurons transiently engage an LHb–RMTg–VTA–NAc coping circuit during stress and are subsequently curtailed through the broader LHA–LHb–RMTg–VTA–NAc–LHA network (Fig. 4i), although the precise mechanisms underlying the suppressive phase remain to be determined.

### Chronic stress impairs D-neuron identity

Having identified an orexin-responsive LHb D-neuron module that promotes active coping, we next asked whether this orexin-dependent regulatory pathway is impaired in depression-like states. To address this, we examined circuit activity and molecular state in two chronic stress models, learned helplessness (LH) and chronic restraint stress (Fig. 5a and Extended Data Fig. 6a–j). In both models, *Fos* expression was increased in LHA orexin neurons and LHb D-neurons. However, this was accompanied by increased *Fos* expression in RMTg GABAergic neurons and reduced *Fos* expression in VTA dopaminergic neurons, with a similar reduction observed in NAc GABAergic neurons (Fig. 5b–f and Extended Data Fig. 6k–o), indicating that, although stress-related upstream recruitment persists, downstream circuit output is reversed under chronic stress.

**Figure 5.**
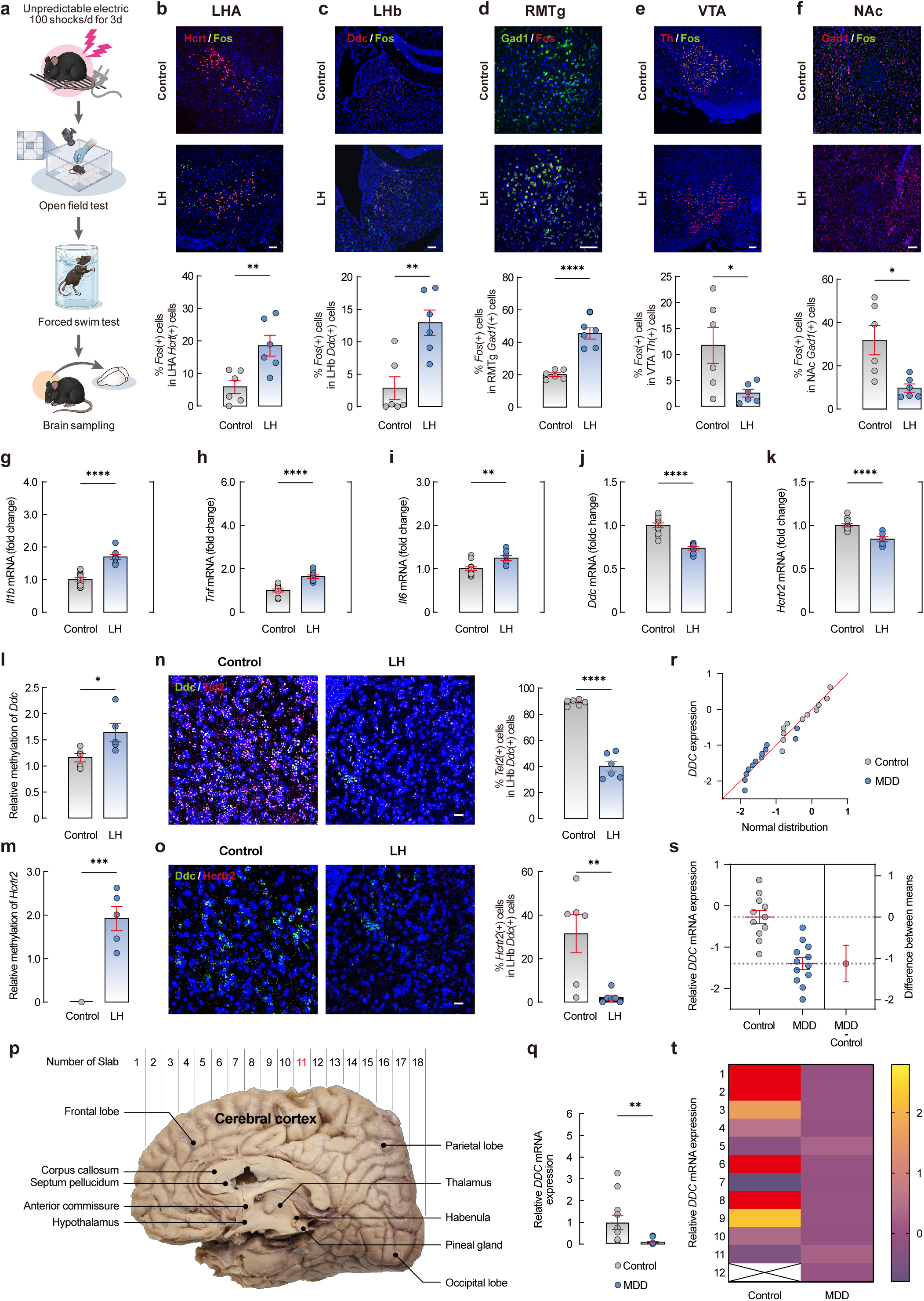
Chronic stress erodes the orexin-responsive LHb D-neuron module. **a**, Experimental schematic of the LH protocol. **b**–**f**, Representative *Fos* images and quantification showing activity in LHA orexin neurons (**b**), LHb D-neurons (**c**), RMTg GABAergic neurons (**d**), VTA dopaminergic neurons (**e**) and NAc GABAergic neurons (**f**) following LH. **g**–**k**, qPCR analysis of LHb transcripts showing expression level of *Il1b* (**g**), *Tnf* (**h**), *Il6* (**i**), *Ddc* (**j**) and *Hcrtr2* (**k**) following LH. **l**,**m**, Promoter methylation analysis of the *Ddc* (**l**) and *Hcrtr2* (**m**) loci in control and LH mice. **n**,**o**, Representative FISH images and quantification showing reduced *Tet2* signal within *Ddc*^+^ neurons (**n**) and reduced *Hcrtr2* signal within *Ddc*^+^ neurons (**o**) following LH. **p**–**t**, Human post-mortem habenula analysis: schematic of the habenula location in coronal slab 11 (**p**), comparison of *DDC* expression between control and MDD cases who died by suicide (**q**), distribution of transformed *DDC* expression values (**r**), group comparison of transformed *DDC* expression by Welch’s *t*-test (**s**), and heat map of human habenular *DDC* expression (**t**). LHA, lateral hypothalamic area; LHb, lateral habenula; RMTg, rostromedial tegmental nucleus; VTA, ventral tegmental area; NAc, nucleus accumbens; LH, learned helplessness; CRS, chronic restraint stress; qPCR, quantitative polymerase chain reaction; FISH, fluorescence *in situ* hybridisation; *Il1b*, interleukin-1β; *Tnf*, tumour necrosis factor; Il6, interleukin-6; *Ddc*, dopa decarboxylase; *Hcrtr2*, hypocretin receptor 2; *Tet2*, ten-eleven translocation 2; MDD, major depressive disorder. Error bars indicate mean ± s.e.m. NS, not significant; \**P* < 0.05, \*\**P* < 0.01, \*\*\**P* < 0.001, \*\*\*\**P* < 0.0001. Scale bars, **b–f**: 100 μm, **n**,**o**: 20 μm. Exact *n* and *P* values are provided in Supplementary Table 5.

Quantitative PCR (qPCR) analysis of the LHb further showed increased cytokine transcripts together with reduced expression of *Ddc* and *Hcrtr2* in both stress models (Fig. 5g–k and Extended Data Fig. 6r–v). This reduction was associated with increased promoter methylation at the *Ddc* and *Hcrtr2* loci. Across animals, methylation levels were inversely correlated with gene expression, consistent with inflammation-associated epigenetic remodelling of the orexin–D-neuron module (Fig. 5l,m and Extended Data Fig. 6e and w–y). FISH analysis further showed reduced *Tet2* and *Hcrtr2* signals within *Ddc*^+^ neurons (Fig. 5n,o), indicating that chronic stress weakens D-neuron identity and orexin responsiveness at the single-cell level, consistent with a circuit-wide shift towards increased LHA–LHb–RMTg recruitment and reduced mesolimbic output (Extended Data Fig. 6z).

In post-mortem human habenula tissue from individuals with major depressive disorder who died by suicide, *DDC* expression was reduced (Fig. 5p–t), whereas *HCRTR2* expression was preserved (Extended Data Fig. 7a–d), suggesting a relative loss of D-neuron identity with preservation of receptor expression. Together, these findings indicate that chronic stress is associated with coordinated inflammatory and epigenetic remodelling in the LHb, accompanied by progressive impairment of D-neuron identity and destabilisation of the orexin-responsive resilience module.

### Orexin-A restores D-neuron function

In light of evidence that orexin-A signalling is altered in some depressive states^36–39^, coupled with our finding that chronic stress is associated with impaired downstream orexin responsiveness in LHb D-neurons, we tested whether replenishing orexin-A in the LHb could rescue LH-induced deficits in LHb D-neurons. To this end, we compared naïve controls with LH-exposed *Ddc*^Cre^ mice receiving intra-LHb vehicle or orexin-A (Extended Data Fig. 8a). Relative to naïve mice, LH markedly reduced active coping behaviour and altered D-neuron-associated transcriptional readouts. Intra-LHb orexin-A significantly increased active coping in the TST compared with LH+vehicle animals and shifted behavioural performance towards naïve-animal levels (Extended Data Fig. 8b). At the molecular level, LH altered selected D-neuron-associated transcriptional readouts, significantly reducing *Ddc* and increasing *Tnf* relative to naïve controls. Intra-LHb orexin-A significantly reversed both changes, whereas *Il1b*, *Il6* and *Hcrtr2* showed no significant group differences (Extended Data Fig. 8c–g). Intra-LHb orexin-A did not significantly alter OFT measures relative to vehicle-treated LH animals, including distance moved, velocity and centre duration, although centre duration remained reduced in LH-exposed groups compared with naïve controls (Extended Data Fig. 8h–j). Together, these findings demonstrate that local restoration of orexin-A signalling is sufficient to reverse stress-induced behavioural and molecular deficits in LHb D-neurons.

We next asked whether this rescue requires OX2R signalling in LHb D-neurons. To address this, we expressed a Cre-dependent short hairpin RNA targeting mouse *Hcrtr2* (shOX2R) or a scrambled control (shScr) in LHb D-neurons. *Hcrtr2* knockdown did not alter total distance moved, mean velocity or centre duration in the OFT, but it increased immobility in the FST (Extended Data Fig. 8k–n), consistent with a pivotal role for endogenous OX2R signalling in mediating active coping behaviour. Following LH, animals were assigned to shScr+vehicle, shScr+orexin-A, shOX2R+vehicle or shOX2R+orexin-A groups (Fig. 6a). Intranasal orexin-A improved active coping in both the TST and FST only in shScr mice; these effects were absent after *Hcrtr2* knockdown, with shOX2R+orexin-A animals remaining comparable to shOX2R+vehicle controls (Fig. 6b,c). Consistent with these behavioural effects, intranasal orexin-A shifted inflammatory and D-neuron-associated transcriptional readouts in a protective direction only in shScr mice, whereas these effects were lost or markedly attenuated after *Hcrtr2* knockdown (Fig. 6d–h). qPCR analysis further confirmed effective *Hcrtr2* knockdown in shOX2R animals (Fig. 6h). Concordantly, orexin-A reduced phosphorylated nuclear factor kappa B p65 signal (p-NF-κB) in mCherry-labelled LHb D-neurons and preserved *Tet2* expression in LHb cells only when OX2R was intact (Fig. 6i–k), without altering OFT locomotor or anxiety-like measures (Fig. 6l–n). These data indicate that the rescue effect of orexin-A depends on OX2R signalling in LHb D-neurons.

**Figure 6.**
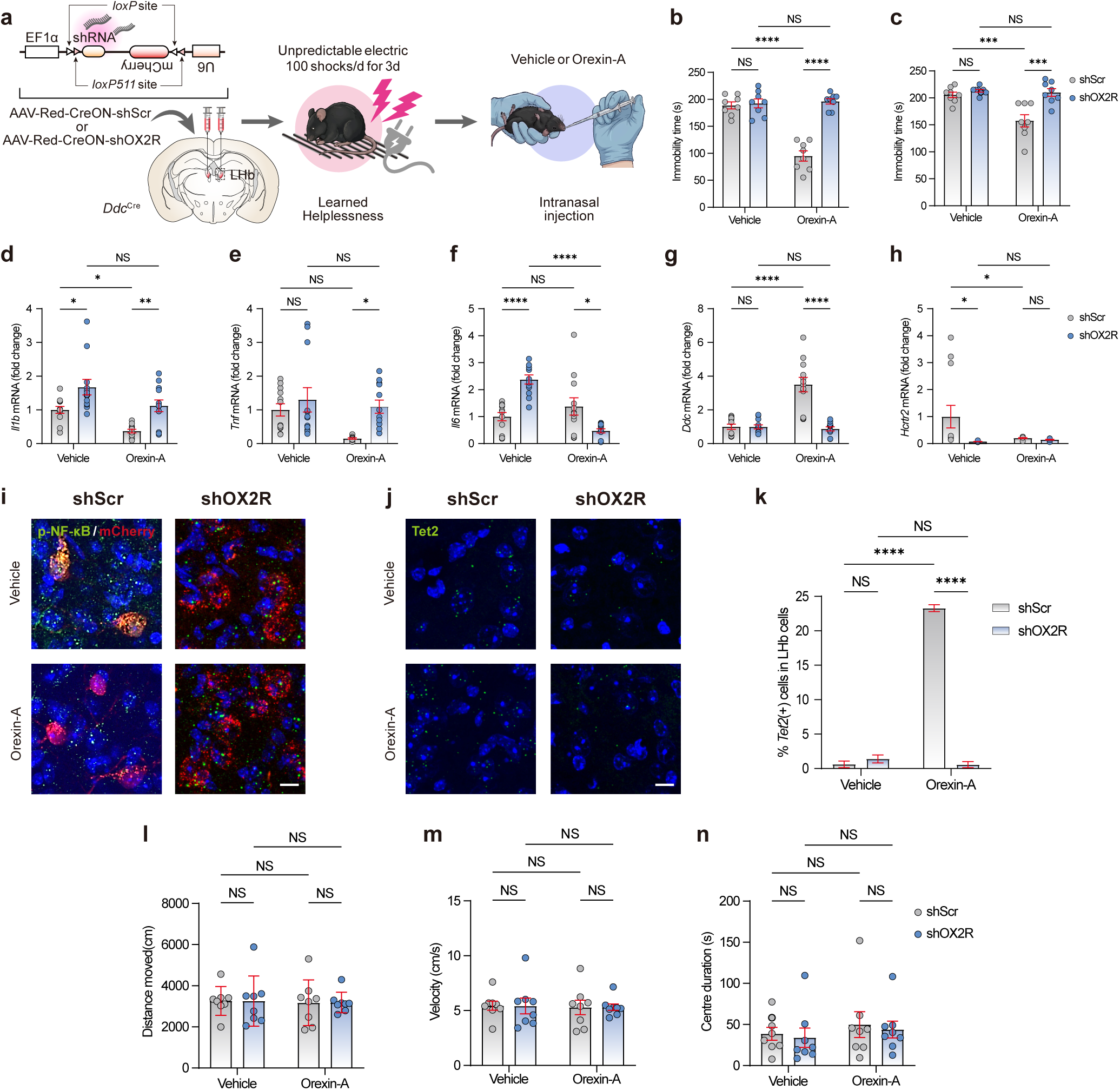
Restoring orexin-A rescues LHb D-neurons through OX2R-dependent inflammatory and epigenetic stabilisation. **a**, Experimental schematic for *Hcrtr2* knockdown in LHb D-neurons using Red-CreON shRNA in *Ddc*^Cre^ mice, followed by LH and intranasal orexin-A administration. **b**,**c**, Quantification of active coping behaviour in the TST (**b**) and FST (**c**), showing that intranasal orexin-A improves coping in shScr mice but not in shOX2R mice. **d**–**h**, qPCR analysis of LHb transcripts showing the effects of intranasal orexin-A and *Hcrtr2* knockdown on *Il1b* (**d**), *Tnf* (**e**), *Il6* (**f**), *Ddc* (**g**) and *Hcrtr2* (**h**). **i**, Representative images showing p-NF-κB (green) in mCherry-labelled LHb D-neurons (red) across conditions. **j**,**k**, Representative images of Tet2 expression in LHb cells (**j**), and quantification of the proportion of *Tet2*^+^ cells in the LHb (**k**), showing preservation by orexin-A in shScr but not in shOX2R mice. **l**–**n**, OFT measures: total distance moved (**l**), mean velocity (**m**), and centre duration (**n**), indicating no significant effect of orexin-A or *Hcrtr2* knockdown on locomotor or anxiety-like behaviour. LHb, lateral habenula; LH, learned helplessness; OX2R, orexin receptor 2; qPCR, quantitative polymerase chain reaction; *Il1b*, interleukin-1β; *Tnf*, tumour necrosis factor; *Il6*, interleukin-6; *Ddc*, dopa decarboxylase; *Hcrtr2*, hypocretin receptor 2; *Tet2*, ten-eleven translocation 2; OFT, open field test; TST, tail suspension test; FST, forced swim test. Error bars indicate mean ± s.e.m. NS, not significant; \**P* < 0.05, \*\**P* < 0.01, \*\*\**P* < 0.001, \*\*\*\**P* < 0.0001. Scale bars, **i**,**j**:10 μm. Exact *n* and *P* values are provided in Supplementary Table 5.

To investigate the intracellular mechanisms linking orexin–OX2R signalling to inflammatory restraint, we next tested whether AMP-activated protein kinase (AMPK) contributes to orexin-dependent rescue. Following LH, *Ddc*^Cre^ mice received intraperitoneal injections of vehicle or the AMPK inhibitor Compound C (CC) 1 h before intranasal administration of orexin-A (Extended Data Fig. 9a). In the OFT, CC pretreatment reduced locomotion, as indicated by decreased total distance moved and mean velocity, while increasing centre duration (Extended Data Fig. 9b–d), indicating that its behavioural effects should be interpreted with caution. Under these conditions, CC pretreatment also increased immobility in both the TST and FST, attenuating the behavioural effects of orexin-A (Extended Data Fig. 9e,f). At the transcriptional level, CC pretreatment did not significantly alter LHb cytokine transcripts or *Hcrtr2* but reduced *Ddc* expression (Extended Data Fig. 9g–k). At the molecular level, orexin-A was associated with low p-NF-κB in mCherry-labelled LHb D-neurons, whereas CC pretreatment increased p-NF-κB under orexin-A treatment (Extended Data Fig. 9l). In parallel, orexin-A preserved *Tet2* expression in LHb cells, and CC pretreatment markedly reduced the proportion of *Tet2*-positive cells in the LHb (Extended Data Fig. 9m,n). Together, these findings support AMPK involvement in orexin-dependent inflammatory restraint and preservation of *Tet2*-associated D-neuron identity, suggesting that the protective actions of orexin-A extend beyond acute behavioural regulation to stabilisation of the *Ddc* programme. Together with prior reports that OX2R triggers Gαq-coupled PLCβ–Ca^2+^ signalling^30,32^, these data are consistent with a model in which orexin-A activates an OX2R–Gαq–PLCβ–Ca^2+^ pathway linked to CaMKKβ-dependent AMPK phosphorylation, restraint of NF-κB activation, preservation of *Tet2*, and demethylation-linked maintenance of the Ddc locus. In this framework, orexin-A is proposed to restore LHb D-neuron function by counteracting stress-associated inflammatory and epigenetic erosion of the D-neuron identity programme.

Collectively, these findings support an OX2R-dependent protective pathway in LHb D-neurons in which orexin-A engages AMPK-linked inflammatory restraint associated with preservation of *Tet2* and maintenance of the *Ddc* programme, thereby restoring stress-impaired D-neuron identity and function.

## Discussion

Our findings identify an orexin-responsive LHb D-neuron module that supports active coping under stress and broadens current views of habenula function. Although the LHb has classically been regarded as an anti-reward hub^8–14^, the present data show that it also contains a molecularly distinct, resilience-associated subpopulation. By engaging OX2R-enriched LHb D-neurons, LHA orexin neurons recruit an LHb–RMTg–VTA–NAc circuit, elevate motivational output and promote active coping. The orexin–habenula interface thus emerges not simply as an accessory branch of canonical aversion circuitry, but as an endogenous mechanism for stabilising motivational output during challenge.

A defining feature of this architecture is that it is self-limiting. LHb-projecting orexin neurons were rapidly recruited at stress onset but were subsequently curtailed after stress, often falling below baseline, consistent with a stress–relief feedback organisation. Considered together with the downstream recruitment pattern and prior evidence for NAc-to-LHA feedback^33–35,40–42^, this biphasic profile suggests that orexin output is mobilised to initiate coping yet actively constrained thereafter to prevent its persistence. In this view, resilience depends not only on mounting an appropriate response to a challenge but also on terminating that response once the challenge has passed. This framework also helps explain why acute stress can robustly recruit the LHb without ordinarily producing depression-like behavioural states: LHb recruitment during challenge can remain adaptive when constrained by feedback and limited in duration.

Our whole-cell patch-clamp recordings in acute brain slices provide a mechanistic framework for how orexin control is implemented in the LHb. Notably, orexin-A and orexin-B exerted distinct effects. These recordings were obtained from LHb neurons rather than genetically identified D-neurons and therefore define orexin responses at the LHb population level; D-neuron specificity is supported by the anatomical enrichment and by the D-neuron-specific OX2R knockdown/rescue experiments. Orexin-A reduced EPSC amplitude in an OX2R-dependent manner and suppressed firing through a Ba^2+^-sensitive inhibitory conductance despite also eliciting a parallel inward current. By contrast, orexin-B did not measurably suppress excitatory synaptic transmission and instead increased spike output while exerting a distinct membrane effect. Together, these findings indicate that orexin signalling in the LHb is ligand-selective rather than uniform, with orexin-A engaging a net inhibitory constraint on neuronal output under the conditions tested.

This interpretation is broadly consistent with prior evidence that intra-LHb orexin-A and orexin-B can produce divergent behavioural effects *in vivo*, although the underlying synaptic mechanisms were not resolved in that study^27^. On the electrophysiological timescale, the Ba^2+^ sensitivity of the orexin-A effect is consistent with recruitment of an inwardly rectifying inhibitory conductance downstream of OX2R signalling. One possible interpretation is that the OX2R-dependent reduction in excitatory synaptic strength reflects recruitment of a Gαi/o-linked pathway, consistent with prior evidence that OX2R can couple to PTX-sensitive G proteins^30–32^. However, because neither cAMP signalling nor NMDAR-isolated synaptic currents were directly examined, the downstream mechanism remains provisional in LHb neurons. In contrast, orexin-B failed to reproduce this inhibitory profile, indicating that its downstream signalling mechanism diverges from that of orexin-A and remains to be fully resolved.

The protective function of this circuit extends beyond acute control of excitability. Restoring orexin-A suppressed NF-κB-associated inflammatory signalling, preserved Tet2, and stabilized the *Ddc* program in an OX2R-dependent manner, whereas these effects were abolished following *Hcrtr2* knockdown in LHb D-neurons. Pharmacological interference with CC attenuated the orexin-A-dependent rescue of molecular readouts and reversed the effects on p-NF-κB and Tet2 preservation, confirming the dependency of the mechanism. Together with prior evidence that OX2R can engage Gαq-coupled PLCβ–Ca^2+^ signalling^30,32^, these results support a model in which orexin-A coordinates rapid control of neuronal output with slower preservation of cellular identity through CaMKKβ–AMPK-linked restraint of NF-κB and Tet2-dependent maintenance of the *Ddc* locus. Because CC also reduced OFT performance, its behavioural effects should be interpreted cautiously; however, the convergent molecular data support AMPK involvement in the protective orexin–OX2R signalling programme.

These results support a working model in which orexin signalling stabilises LHb D-neuron function across multiple levels, from circuit dynamics to intracellular inflammatory and epigenetic control (Extended Data Fig. 10). In this framework, LHA-derived glutamate and orexin jointly engage LHb D-neurons^27,43,44^, which in turn suppress the RMTg and promote downstream mesolimbic output, while a feedback loop through the broader LHA–LHb–RMTg–VTA–NAc–LHA network constrains the duration of this state^5,10,33–35,40–43,45^. At the cellular level, orexin-A appears to recruit both an acute inhibitory branch, compatible with recruitment of an OX2R-linked inwardly rectifying inhibitory conductance, and a slower protective branch, compatible with Gαq–Ca^2+^–CaMKKβ–AMPK-mediated restraint of NF-κB, preservation of *Tet2* and maintenance of *Ddc* expression. The same model also provides a conceptual basis for the orexin-A versus orexin-B dissociation, with orexin-A favouring a self-limiting protective programme and orexin-B favouring increased output without reproducing the inhibitory phenotype.

Chronic stress dismantled this resilience module at multiple levels. Across depression-like models, orexin-related upstream recruitment persisted or increased, yet adaptive downstream output was reversed; meanwhile, *Ddc* and *Hcrtr2* expression, Tet2 signalling, and D-neuron identity were progressively eroded in parallel with inflammatory induction and promoter methylation. Thus, the chronic stress state is best interpreted not as a failure of a single component, but as a progressive collapse of a multi-layered module involving inflammatory activation, epigenetic remodelling, loss of D-neuron identity, and impaired orexin responsiveness. These convergent changes suggest that prolonged stress does not simply silence the system; rather, it uncouples stress-driven recruitment from adaptive output by eroding both D-neuron identity and orexin responsiveness. The reduction of *DDC*, with relative preservation of *HCRTR2*, in post-mortem human habenula samples further supports the translational relevance of this framework. In this context, impaired orexin-A signalling efficacy within the LHb could leave neuronal excitability less effectively restrained and potentially shift orexin influence towards a more orexin-B-like excitatory state, thereby favouring LHb hyperactivity.^27,36–39,46^.

Importantly, this stress-induced state remained reversible. Replenishing orexin-A restored active coping and reversed molecular deficits in an OX2R-dependent manner. This rescue profile suggests that chronic stress does not eliminate the module entirely, but instead drives it into a destabilised state that remains amenable to reactivation upon restoration of the appropriate signalling pathway. Notably, this endogenous orexin-A pathway may converge functionally with ketamine-sensitive mechanisms in modulating LHb output; both appear capable of constraining LHb hyperactivity and reinstating adaptive behavioural responsiveness^11,47,48^. At a broader systems level, this convergence may also extend to stress-associated inflammatory signalling, although current evidence for ketamine’s anti-inflammatory effects remains primarily systemic rather than LHb-specific^49,50^.

Several limitations should be noted. The behavioural effects of CC should be interpreted with caution given its suppression of open-field locomotion. Although our rescue experiments implicate AMPK, they do not establish whether CC acts directly on LHb D-neurons or via broader circuit-level effects. Furthermore, as the biphasic dynamics were recorded from LHb-projecting LHA orexin neurons, they primarily support presynaptic or circuit-level regulation; nonetheless, postsynaptic adaptation of OX2R signalling remains a plausible additional mechanism. The post-mortem human findings are associative and cannot exclude contributions from medication history or other confounders. Finally, while our data demonstrate an OX2R-dependent role for orexin in LHb D-neurons, LHA orexin neurons co-release glutamate, and a substantial non-D-neuron *Hcrtr2*^+^ population exists within the LHb; future studies are required to resolve the relative contributions of these components.

More broadly, these findings define a self-limiting orexin–habenula resilience circuit that links stress coping to inflammatory and epigenetic homeostasis. The collapse of such intrinsically self-limiting modules may therefore represent a general mechanism by which chronic stress is converted into persistent motivational dysfunction.

## Methods

### Animals

Adult C57BL/6J, *Orexin*^Cre^ and *Ddc*^Cre^ mice aged 8–12 weeks were used in this study. Unless otherwise indicated, male mice were used. Mice were group-housed under a 12-h light/dark cycle with ad libitum access to food and water. All procedures were approved by the Korea University Institutional Animal Care and Use Committee (KOREA-2023-0135). Detailed animal information is provided in the Supplementary Methods.

### Viral vectors, stereotaxic surgery and circuit manipulations

Adeno-associated viral vectors were packaged in-house using serotype 2/9 capsids. Stereotaxic injections, fibre implantation and guide cannulation were performed under isoflurane anaesthesia using established coordinates for the LHb, LHA and NAc. For monosynaptic retrograde tracing, Cre recombinase (Cre)-dependent helper viruses were injected into the LHb, followed by EnvA-pseudotyped glycoprotein-deleted rabies virus. For anterograde tracing, a Cre-dependent synaptophysin-labelled reporter was injected into the LHA. Optogenetic, chemogenetic, knockdown and dopamine-sensor experiments were performed using appropriate Cre- or flippase recombinase (FLPo)-dependent viral strategies. Detailed plasmid information, viral constructs, titres and surgical procedures are provided in the Supplementary Methods.

### Behavioural assays and stress models

Behavioural assays included the OFT, TST, FST, CPP and RTPP. Acute stress was induced by tail suspension, chronic stress by chronic restraint stress, and learned helplessness by repeated inescapable footshocks. Optogenetic stimulation was applied during OFT, TST and RTPP, whereas chemogenetic activation was assessed in OFT, TST, FST and CPP. Full behavioural procedures, stimulation schedules and stress protocols are described in the Supplementary Methods.

### Chemogenetics and optogenetics

To manipulate LHb-projecting orexin neurons, *Orexin*^Cre^ mice were subjected to intersectional viral strategies targeting the LHb and LHA. For chemogenetic activation, LHb-projecting orexin neurons expressing hM3Dq were activated by systemic CNO administration. For optogenetic experiments, ChR2-expressing orexin projections were stimulated with 473-nm light through bilaterally implanted optical fibres above the LHb. In a separate cohort, nucleus accumbens dopamine dynamics were monitored using the GRAB-rDA3m sensor. Detailed viral strategies, stimulation parameters, CNO administration and fibre photometry procedures are provided in the Supplementary Methods.

### Electrophysiology

*In vivo* single-unit recordings were performed using optrodes implanted above the LHA to identify LHb-projecting orexin neurons by optotagging during acute stress. *In vitro* whole-cell recordings were performed in acute coronal slices containing the LHb to assess spontaneous synaptic transmission, membrane currents, firing responses and orexin receptor pharmacology. Light-evoked responses were measured in slices from mice expressing ChR2 in LHA orexin neurons. Detailed recording conditions, solutions, inclusion criteria and analysis procedures are provided in the Supplementary Methods.

### Histology and molecular analyses

FISH, immunohistochemistry, qPCR and DNA methylation analyses were used to quantify molecular changes across the LHA, LHb, RMTg, VTA and NAc. For rescue experiments, OX2R was activated either by intra-LHb orexin-A infusion or by intranasal orexin-A administration, with or without CC pretreatment. Human habenular *DDC* and *HCRTR2* expression was measured in archived post-mortem tissue from a previously characterised cohort. Detailed probe, antibody, primer and assay information is provided in the Supplementary Methods and Supplementary Tables.

### Statistics and reproducibility

Data are presented as mean ± s.e.m. Statistical analyses were performed using GraphPad Prism 10 (GraphPad Software, Boston, MA, USA), unless otherwise stated. Normality was assessed using the Shapiro–Wilk test. Depending on the experimental design, two-tailed unpaired or paired Student’s *t*-tests, one-way ANOVA, repeated-measures one-way ANOVA, or two-way ANOVA were used with appropriate *post hoc* multiple-comparisons tests. Exact statistical tests, sample sizes, and exact *P* values are provided in the figure legends and Supplementary Table 5. Mice were randomly assigned to experimental groups, and data acquisition and analysis were performed blind to group allocation whenever feasible. Data were excluded only on the basis of pre-established technical criteria described in the Supplementary Methods. Sample sizes were estimated on the basis of prior literature, preliminary effect sizes and feasibility; no formal statistical methods were used to predetermine sample size.

## Data availability

All data supporting the findings of this study are available within the paper and its Supplementary Information. Source data for all main and extended data figures are provided with this paper. Additional datasets generated and/or analysed during the current study are available from the corresponding author upon reasonable request.

## Acknowledgements

We thank Professor Takeshi Sakurai (International Institute for Integrative Sleep Medicine [WPI-IIIS], University of Tsukuba, Japan) for kindly providing the *Orexin*^Cre^ mice. We thank Professor Kihoon Han (Korea University College of Medicine), Professor Gi Hoon Son (Korea University College of Medicine), and Professor Jee Hoon Roh (Korea University College of Medicine) for their critical review of this paper and their feedback. This work was supported by the National Research Foundation of Korea (NRF) grants funded by the Korea government (MSIT) (NRF-2022R1A6A3A01087533 and RS-2024-00340080 to S.H.Y.; RS-2023-00272290 to H.K.).

## Contributions

S.H.Y. and H.W.L. conceived the study. S.H.Y. designed the study, performed surgical procedures, viral tracing, and data analysis. S.H.Y. and J.T.J. performed optogenetic, chemogenetic, and pharmacological experiments and established the learned helplessness model. E.Y. performed FISH and histological analyses. J.L. and S.C. carried out whole-cell patch-clamp recordings. G.H.P. and Y.S.J. performed optrode recordings and fibre photometry. H.S.P. produced recombinant AAVs, performed qPCR, and analysed human habenular *DDC* and *HCRTR2* expression. J.W.M. carried out DNA methylation analyses. J.–Y.P. generated AAV vectors for Cre-dependent *Hcrtr2* shRNA and scrambled controls. K.–J.B. provided the chronic restraint stress model. D.L. and H.Y. analysed the data and prepared the figures. H.K. provided overall supervision. S.H.Y. wrote the manuscript with input from all authors.

