## Supplementary Information, Source Data, and Supplementary Table 5 for "A self-limiting orexin–habenula circuit for stress resilience": Yang-et-al-Nature-Supplementary-Information.pdf

### Supplementary Methods

#### Animals

Adult mice aged 8–12 weeks were used in this study. Wild-type C57BL/6J mice were purchased from Dae Han Bio Link (DBL, Eumseong, Republic of Korea). *Orexin*$^{\mathrm{Cre}}$ mice were kindly provided by Professor Takeshi Sakurai (International Institute for Integrative Sleep Medicine [WPI-IIIS], University of Tsukuba, Tsukuba, Japan), and *Ddc*$^{\mathrm{Cre}}$ mice (B6.FVB(Cg)-Tg(Ddc-cre)SD56Gsat/Mmucd; RRID:MMRRC_037410-UCD) were obtained from the Mutant Mouse Resource and Research Center (MMRRC, University of California, Davis, CA, USA). Unless otherwise indicated, male mice were used to minimise potential behavioural variability associated with the oestrous cycle. Mice were group-housed (3–4 per cage) in temperature- and humidity-controlled rooms under a 12-h light/dark cycle (lights on at 08:00), with ad libitum access to food and water. All animals were acclimatised to the animal facility for at least 1 week before any procedures. All experimental procedures were approved by the Korea University Institutional Animal Care and Use Committee (IACUC; approval no. KOREA-2023-0135) and were conducted in accordance with institutional and national guidelines.

#### Viruses

Adeno-associated viruses (AAVs) with serotype 2/9, 8, and 5 capsids were packaged in HEK293 cells and purified by iodixanol density gradient ultracentrifugation. Viral titres were determined by real-time PCR using ITR-specific primers, with standard curves generated from plasmid DNA containing the ITR sequence. Purified AAV stocks were aliquoted to minimize freeze–thaw cycles and stored at –80 °C at a concentration of approximately $1 \times 10^{13}$ vg/ml until use.

The following plasmids were obtained from Addgene (Watertown, MA, USA): pAAV-hSyn-DIO-hM3D(Gq)-mCherry (#44361), pAAV-hSyn-DIO-mCherry (#50459), pAAV-hSyn-GRAB-rDA3m (#208702), pAAV-hSyn-FLEX-mGFP-2A-synaptophysin-mRuby (#71760), pAAV-EF1α-DIO-hChR2(H134R)-EYFP-WPRE-HGHpA (#20298), pAAV-EF1α-fDIO-hChR2(H134R)-EYFP (#55639), pAAV-hSyn-fDIO-hM3D(Gq)-mCherry-WPREpA (#154868), pAAV-hSyn-fDIO-mCherry-WPREpA (#177328), pAAV-EF1α-DIO-FLPo-WPRE-hGHpA (#87306), CAG-FLEX-TCB (#48333), CAG-FLEX-RG (#48332), and rAAV2-retro helper (#81070). These plasmids were used as transfer or helper constructs for AAV production, as indicated in the relevant experimental sections.

For monosynaptic tracing experiments, EnvA-pseudotyped, glycoprotein-deleted rabies virus expressing GFP (RVΔG-GFP) was obtained from the Viral Vector Core at the Kavli Institute for Systems Neuroscience, Norwegian University of Science and Technology (NTNU, Trondheim, Norway). Viral aliquots were stored at −80 °C and thawed immediately before stereotaxic injection into the target regions.

For OX2R knockdown, Cre recombinase (Cre)-dependent AAV-shRNA vectors were generated using pAAV-Red-CreON-shRNA [Control], a gift from Eun Mi Hwang (Addgene #180775). In this system, shRNA is expressed from the U6 promoter and mCherry from the EF1α promoter following Cre-dependent recombination. A 21-nucleotide sequence targeting mouse OX2R was designed based on the same cDNA template (#MG50862, Sino Biological, Beijing, China) and inserted downstream of the U6 promoter. A scrambled shRNA sequence with no predicted targets in the mouse genome was inserted into the same backbone to generate a control vector with identical architecture. Both knockdown and control viruses were packaged in-house, and viral titres were determined by real-time PCR.

**Stereotaxic surgery and viral delivery**

Mice were anaesthetised with isoflurane (5% for induction and 1% for maintenance) and placed in a stereotaxic frame (Ultra Precise Mouse Stereotaxic Instruments; Stoelting Co., Wood Dale, IL, USA). Following a midline scalp incision, a small craniotomy was made using a handheld drill. Viral vectors were delivered through a 30-gauge microinjection cannula (P1 Technologies, Roanoke, VA, USA) connected to an UltraMicroPump III (World Precision Instruments, Sarasota, FL, USA) at a rate of 50 nL $min^{-1}$. After injection, the scalp was closed with 9-mm autoclips (#205016, MikRon Precision, Inc., Garfield, NJ, USA), and mice received post-operative antibiotics and analgesics. Animals were then transferred to a clean cage placed on a heating pad and monitored until full recovery from anaesthesia. Mice were returned to their home cages and allowed to recover for 3 weeks before behavioural or physiological experiments to ensure sufficient viral expression.

Stereotaxic coordinates were defined relative to bregma as follows: lateral habenula (LHb), anterior–posterior (AP) −1.58 mm, medial–lateral (ML) ±0.90 mm, dorsal–ventral (DV) −3.10 mm, with a 10° mediolateral approach angle; lateral hypothalamus (LHA), AP −1.10 mm, ML ±1.60 mm, DV −4.58 mm, with a 10° mediolateral approach angle; and nucleus accumbens (NAc), AP +1.20 mm, ML +1.60 mm, DV −4.00 mm. For pharmacological microinfusion into the LHb, guide cannulae were implanted at AP −1.78 mm, ML −0.50 mm and DV −2.30 mm from the skull surface using a vertical approach.

### Circuit tracing

For monosynaptic retrograde tracing, *Ddc*$^{Cre}$ mice received a 1:1 mixture of AAV5-CAG-FLEX-TCB (TVA-mCherry receptor) and AAV8-CAG-FLEX-RG (rabies glycoprotein) into the LHb to enable Cre-dependent expression of the components required for RVΔG infection and trans-synaptic spread. After 2 weeks, RVΔG-GFP was injected into the same LHb coordinates. Mice were perfused 1 week later for histological analysis.

For anterograde tracing of orexin projections, *Orexin*$^{Cre}$ mice received AAV2/9-hSyn-FLEX-mGFP-2A-synaptophysin-mRuby into the LHA to label orexin neuron somata and axon terminals. Mice were allowed to recover for 3 weeks before brain collection and fluorescence imaging. Orexin projections to LHb D-neurons were assessed by combining anatomical tracing with fluorescence *in situ* hybridisation.

### Behavioural assays

All behavioural experiments were conducted during the light phase in a temperature- and humidity-controlled room. Mice were habituated to the testing room for at least 30 min before each session. Apparatuses were cleaned with 70% ethanol between trials. Behaviour was recorded and analysed using EthoVision XT 12 (Noldus Information Technology BV, Wageningen, the Netherlands; RRID: SCR_000441), unless otherwise stated.

### Open field test (OFT)

The OFT was used to assess general locomotor activity and anxiety-like behaviour. Mice were placed in the centre of a white square arena (45 × 45 × 40 cm) and allowed to explore freely. Total distance travelled, mean velocity and time spent in the centre zone were quantified automatically.

**Tail suspension test (TST)**

The TST was used to assess passive coping behaviour. Mice were suspended by the tail using adhesive tape placed approximately 1 cm from the tip and positioned 29 cm above the floor. Sessions were video-recorded, and immobility during the final 4 min was scored manually at 5-s intervals by an experimenter blinded to condition. Immobility was defined as the complete absence of limb movement. Total immobility time was calculated by multiplying the number of immobile epochs by 5 s.

**Forced swim test (FST)**

The FST was used to assess passive coping behaviour. Mice were placed individually in a transparent glass cylinder (23.5 cm in height, 13 cm in diameter) filled with water (23–25 °C) to a depth of 17 cm. Each session lasted 6 min, and immobility during the final 4 min was scored using the same criteria as for the TST. Water was changed between animals to minimise olfactory confounds.

**Conditioned place preference test (CPP)**

CPP was used to assess the reinforcing effects of chemogenetic manipulation. The apparatus consisted of three chambers with distinct visual and tactile cues. On day 1 (pre-test), mice were allowed to explore all chambers freely for 15 min. On conditioning days, mice were confined to one side chamber after vehicle treatment and to the opposite chamber after Clozapine-N-oxide (CNO; BML-NS105, Enzo Life Sciences, Farmingdale, NY, USA) treatment, with chamber assignment counterbalanced across animals. On day 8 (post-test), mice were again allowed to explore all chambers freely for 15 min. Preference scores were calculated as the change in time spent in the CNO-paired chamber relative to baseline.

**Real-time place preference test (RTPP)**

RTPP was used to assess the reinforcing effects of optogenetic stimulation. The apparatus consisted of three connected chambers with distinct visual and tactile cues. Mice were placed in the central chamber and allowed to explore freely for 20 min. Time spent in each chamber, distance travelled and mean velocity were recorded automatically. Stimulation contingency is described in the Optogenetics section below.

**Chemogenetics**

To selectively activate orexin projections from the LHA to the LHb, *Orexin*$^{Cre}$ mice received stereotaxic injections of rAAV2-EF1α-DIO-FLPo into the LHb and either AAV2/9-hSyn-fDIO-hM3D(Gq)-mCherry or control AAV2/9-hSyn-fDIO-mCherry into the LHA. This dual-recombinase strategy enabled flippase recombinase (FLPo)-dependent expression of hM3D(Gq) specifically in LHb-projecting orexin neurons.

In a separate cohort used for dopamine monitoring, mice additionally received AAV2/9-hSyn-GRAB-rDA3m into the NAc, and an optical fibre was implanted 200–300 μm above the viral injection site. CNO was dissolved in sterile 1× PBS and administered intraperitoneally at 1 mg $kg^{-1}$, 40 min before recording or behavioural testing.

Behavioural testing was performed using OFT, TST, FST and CPP, as described above. In the dopamine-monitoring cohort, NAc dopamine dynamics were recorded by fibre photometry following CNO administration to assess mesolimbic responses to chemogenetic activation of LHb-projecting orexin neurons. Detailed fibre photometry procedures are described below.

Following behavioural assessments or dopamine recording, mice were perfused, and brains were collected for post hoc molecular analysis. Coronal brain sections (14 μm) were prepared and processed for FISH to quantify *Fos* expression and region-specific molecular markers within the LHA, LHb, rostromedial tegmental area (RMTg), ventral tegmental area (VTA) and NAc.

**Fibre photometry**

Dopamine signals were recorded using Synapse software and a Tucker-Davis Technologies processor. Optical signals were generated using a fluorescence Mini Cube (Doric Lenses, Quebec, QC, Canada) coupled to sinusoidally modulated LEDs at 465 nm (531 Hz) and 405 nm (211 Hz). Excitation light was delivered through a low-autofluorescence fibre-optic patch cord (400 μm core diameter, numerical aperture (NA) 0.48; Doric Lenses), and emitted fluorescence was collected and detected using a photoreceiver (LxPS1, Tucker-Davis Technologies, Alachua, FL, USA). Light power was calibrated at the fibre tip and maintained at 30 μW throughout the recording sessions. The 405 nm signal was used as an isosbestic control to correct for motion artefacts and photobleaching. ΔF/F was calculated as $(F_{465} - F_{405\ fitted}) / F_{405\ fitted}$, and the resulting ΔF/F signal was normalised to z-scores using the baseline period (−5 to −3 s relative to event onset). Dopamine signals were aligned to sucrose lick onset (time 0), and peri-event activity was analysed over a −5 to +13 s window using custom MATLAB® (R2023b, The MathWorks, Natick, MA, USA) scripts. For quantification, the area under the curve (AUC) of the z-scored ΔF/F trace was calculated for predefined time windows relative to sucrose lick onset (baseline: −5 to −3 s; post-event: 0–0.2 s and 1–5 s).

**Optogenetics**

To selectively activate orexin projections to the LHb, *Orexin*$^{Cre}$ mice received stereotaxic injections of AAV2/9-EF1α-DIO-hChR2(H134R)-EYFP into the LHA. Optical fibres (200-μm core diameter, NA 0.37; Thorlabs, Newton, NJ, USA) were implanted bilaterally 200–300 μm above the LHb. Mice were allowed to recover for 3 weeks to ensure robust viral expression before behavioural testing. Behavioural testing was performed using OFT, TST and RTPP, as described above. During stimulation, 473-nm blue light was delivered from a 473-nm blue DPSS laser system (BL473T2-100 mW; Shanghai Laser & Optics Century Co., Ltd., Shanghai, China) through patch cords connected to the implanted fibres. Stimulation consisted of 5-ms pulses at 30 Hz, delivered at 5–10 mW measured at the fibre tip. In the OFT, mice underwent alternating 3-min laser-OFF and 3-min laser-ON epochs. In the TST, mice underwent a 3-min laser-OFF epoch followed by a 3-min laser-ON epoch and a final 3-min laser-OFF epoch. In the RTPP assay, optical stimulation was delivered only while the animal occupied the chamber paired with photostimulation and was terminated upon exit from that chamber. At the end of the experiments, mice were transcardially perfused, and brains were collected. Coronal brain sections (40 μm) were prepared using a cryostat, and viral expression and fibre placement were verified by native EYFP fluorescence. Animals were excluded if *post hoc* histological analysis showed mistargeted viral expression or incorrect fibre placement relative to the LHb.

***In vivo* electrophysiology**

To selectively target LHb-projecting orexin neurons, *Orexin*$^{Cre}$ mice received stereotaxic injections of rAAV2-EF1α-DIO-FLPo into the LHb and AAV2/9-EF1α-fDIO-hChR2(H134R)-EYFP into the LHA. An optrode array consisting of a 200-μm optic fibre (NA 0.22, Thorlabs) and four 20-μm tungsten tetrodes (California Fine Wire Co., Grover Beach, CA, USA) was implanted above the LHA.

Following recovery, mice were connected to a 473-nm diode-pumped laser (Laserglow Technologies, Toronto, ON, Canada) and a Digital Lynx acquisition system (Neuralynx, Bozeman, MT, USA). Neural signals were amplified 1,000–8,000×, filtered between 0.6 and 6 kHz, and digitised at 32 kHz. During optotagging, trains of ten blue-light pulses (5-ms pulse width, 30 Hz, 4–12 mW $mm^{-2}$) were delivered through the optic fibre. If no light-responsive units were detected, the optrode was advanced in 40–80 μm increments, to a maximum of 160 μm per day. Behavioural recording commenced on the day after at least two light-responsive units had been identified.

Each recording session consisted of a 15-min baseline period in a holding cage, 10-min of tail suspension stress, and 24-min of post-stress observation. At the end of each session, ten trains of ten light pulses were delivered to reconfirm light-responsive units. In total, 30 recording sessions were obtained from five mice (5–7 sessions per animal).

Spikes were sorted offline on the basis of waveform features using Offline Sorter™ (Plexon Inc., Dallas, TX, USA). Only units with stable firing throughout the recording period were included in the analysis, which was performed in MATLAB. Orexin neurons were defined as units showing a spike probability greater than 0.7 and a response latency shorter than 5.7 ms across 100-light pulses, together with a high waveform correlation between spontaneous and light-evoked spikes.

#### *In vitro* electrophysiology

Acute coronal brain slices (300 μm thick) containing the LHb were prepared from mice deeply anaesthetised with isoflurane and rapidly decapitated. Brains were quickly removed and sectioned in ice-cold, oxygenated N-methyl-D-glucamine (NMDG)-based cutting solution containing (in mM): 92 NMDG, 2.5 KCl, 1.25 $NaH_2PO_4$, 30 $NaHCO_3$, 20 HEPES, 0.5 $CaCl_2$, 10 $MgCl_2$ and 25 glucose (pH 7.3–7.4, adjusted with HCl; 300–310 mOsm), continuously bubbled with 95% $O_2$/5% $CO_2$. Slices were then transferred to an oxygenated recovery solution containing (in mM): 92 NaCl, 2.5 KCl, 1.25 $NaH_2PO_4$, 30 $NaHCO_3$, 20 HEPES, 2 $CaCl_2$, 2 $MgCl_2$ and 25 glucose, incubated at 30 °C for 60 min, and subsequently maintained at room temperature (22–24 °C) for at least 30 min before recording. During recordings, slices were continuously perfused at 1.5–2 mL min−1 with oxygenated artificial cerebrospinal fluid (ACSF) containing (in mM): 124 NaCl, 2.5 KCl, 1.25 $NaH_2PO_4$, 24 $NaHCO_3$, 1.5 $CaCl_2$, 1.5 $MgCl_2$ and 10 glucose.

LHb neurons were visualised using differential interference contrast optics on an upright microscope (BX51WI, Olympus, Tokyo, Japan). Whole-cell patch-clamp recordings were obtained using borosilicate glass pipettes (3–5 MΩ) filled with an internal solution containing (in mM): 135 K-gluconate, 10 HEPES, 0.2 EGTA, 2 $MgCl_2$, 4 Mg-ATP and 0.3 Na-GTP (pH 7.3, adjusted with KOH; 290–300 mOsm). Signals were amplified using a MultiClamp 700B amplifier (Molecular Devices, San Jose, CA, USA), low-pass filtered at 2 kHz, and digitised at 10 kHz using pClamp software (v10.x, Molecular Devices). Series resistance was monitored throughout the recordings; cells with an initial series resistance >20 MΩ or a change exceeding 20% during recording were excluded from analysis. Recordings were also excluded if the holding current varied by >50 pA from baseline. The liquid junction potential (~10 mV) was not corrected.

Spontaneous excitatory postsynaptic currents (sEPSCs) were recorded in voltage-clamp mode at a holding potential of −60 mV in the presence of bicuculline (10 μM) and picrotoxin (50 μM) to block $GABA_A$ receptor-mediated currents.

For optogenetic experiments, *Orexin*$^{Cre}$ mice received stereotaxic injections of AAV2/9-EF1α-DIO-ChR2-EYFP into the LHA. ChR2-EYFP expression in orexin neurons was confirmed by epifluorescence imaging before electrophysiological recording. Simultaneous patch-clamp recording and optical stimulation were performed using an Optopatcher™ (A-M Systems, Sequim, WA, USA), which enabled light delivery through the patch pipette while maintaining stable whole-cell access. Light-evoked synaptic responses were elicited by 473-nm blue light pulses (5-ms, 30 Hz) delivered using an LED-based optogenetic stimulation system (CoolLED, Andover, UK) coupled to the Optopatcher, thereby providing spatially restricted illumination of ChR2-expressing axonal projections in the vicinity of the recorded neuron. Light intensity was calibrated to approximately 5 mW mm$^{-2}$ at the specimen plane, and light delivery was controlled by TTL-triggered pulses synchronised with electrophysiological data acquisition.

Pharmacological effects of orexin receptor signalling were assessed by bath application of orexin-A (500 nM; #1455, Tocris Bioscience, Bristol, UK), orexin-B (500 nM; #1457, Tocris Bioscience) or the selective OX2R antagonist TCS-OX2-29 (10 μM; #3371, Tocris Bioscience) for 5–10 min. To assess the contribution of $Ba^{2+}$-sensitive inwardly rectifying $K^+$ conductances, cells were held at −60 mV and a series of hyperpolarising voltage steps (−100 to −60 mV, in 10-mV increments, 500-ms duration) was applied before and after bath application of orexin-A (500 nM).

Spontaneous action potential firing was recorded in current-clamp mode without current injection. Firing rates were quantified over 60–120 s epochs under baseline conditions and following bath application of orexin-A, with or without subsequent application of $BaCl_2$. Membrane potential was monitored throughout. Cells exhibiting unstable baseline firing or membrane potential drift exceeding ±3 mV were excluded from analysis.

All slice recordings were performed at room temperature (22–24 °C) unless otherwise stated. Experimenters were blinded to treatment group, and slices were randomly assigned to recording conditions. Data are reported as the number of cells (n) obtained from at least three independent animals per group. Key findings were replicated across independent experimental cohorts.

**Stress models**

Acute stress was induced by a 10-min tail suspension procedure. After at least 1 week of habituation to the home cage environment, mice were suspended by the tail for 10 min to elicit acute stress responses.

For the chronic restraint stress (CRS) model, mice were individually placed in well-ventilated 50-mL polypropylene conical tubes (#54050, SPL Life Sciences, Pocheon, Republic of Korea) for 2 h per day over 20 consecutive days. Control mice remained in their home cages without restraint.

For the learned helplessness (LH) model, mice were placed in conditioning chambers (30 × 30 × 25 cm; Multi Conditioning System, TSE Systems, Chesterfield, MO, USA) and subjected to 100 inescapable footshocks (0.3 mA, 5-s duration) delivered at pseudorandom intershock intervals ranging from 5 to 99 s. Control mice were placed in the same chambers for an equivalent period but did not receive footshocks.

**Fluorescent *in situ* hybridisation (FISH)**

FISH was performed to quantify mRNA expression across multiple brain regions, including the LHA, LHb, RMTg, VTA and NAc. Brains were rapidly removed and flash-frozen on dry ice. Coronal sections (14 μm thick) were prepared using a CM3050 S cryostat (Leica Microsystems,

Wetzlar, Germany) and mounted on Superfrost Plus™ microscope slides (Thermo Fisher Scientific, Waltham, MA, USA).

*In situ* hybridisation was performed using the RNAscope™ Multiplex Fluorescent V2 Assay (Advanced Cell Diagnostics, Newark, CA, USA) according to the manufacturer's instructions. Briefly, sections were fixed in 4% paraformaldehyde for 10 min at 4 °C, dehydrated through a graded ethanol series and treated with RNAscope™ Protease IV. Target-specific probes (Supplementary Table 1) were hybridised at 40 °C in a HybEZ™ II oven (Advanced Cell Diagnostics). Signal amplification and detection were performed using TSA Vivid™ Fluorophores (Vivid 520/570/650). Sections were counterstained with DAPI to visualise nuclei.

Slides were coverslipped with ProLong™ Gold Antifade Mountant (Invitrogen, Thermo Fisher Scientific, Carlsbad, CA, USA). Images were acquired using a Leica TCS SP8 confocal laser-scanning microscope (Leica Microsystems) equipped with a 40× oil-immersion objective (NA 1.3). Images were acquired and processed using LAS X software (Leica Application Suite X, Leica Microsystems), with laser power, gain and offset held constant across experimental groups.

Quantification was performed using HALO® software (Indica Labs, Albuquerque, NM, USA) and ImageJ (NIH, Bethesda, MD, USA). For each brain region, at least three non-overlapping fields and a minimum of 50 cells per mouse were analysed. All analyses were performed blind to experimental group allocation. Data are presented as the mean number of RNA puncta per cell and puncta density per unit area. Probe information is provided in Supplementary Table 1.

**Immunohistochemistry (IHC)**

Animals were deeply anaesthetised with a mixture alfaxalone (40 mg $kg^{-1}$) : xylazine (10 mg $kg^{-1}$) and transcardially perfused with ice-cold phosphate-buffered saline (PBS), followed by 4% paraformaldehyde (PFA) in PBS. Brains were removed, post-fixed in 4% PFA overnight at 4 °C, and cryoprotected in 30% sucrose for at least 48 h until they sank. Coronal sections (40 µm thick) were prepared using a CM3050 S cryostat (Leica Microsystems) and collected from multiple brain regions, including the MS, LS, LPO, LHA and LHb.

Free-floating sections were incubated for 1 h at 37 °C in PBS containing 3% bovine serum albumin and 0.2% Triton X-100, followed by incubation with primary antibodies diluted in the same buffer. The following primary antibodies were used: rabbit anti-orexin-A (1:500; #ab6214, Abcam, Cambridge, UK), chicken anti-GFP (1:1,000; #ab13970, Abcam) and rabbit anti-NF-κB p65 (phospho-Ser536) (1:500; #3033, Cell Signaling Technology, Danvers, MA, USA).

After washing in PBS, sections were incubated for 2 h at room temperature with species-appropriate secondary antibodies diluted in the same buffer. Nuclei were counterstained with Hoechst 33342 (1:1,000; #H3570, Thermo Fisher Scientific) for 10 min at room temperature. Sections were mounted and imaged using a Leica TCS SP8 confocal microscope (Leica Microsystems).

**Quantitative PCR (qPCR)**

qPCR was performed in according to established protocols. Following euthanasia, Mouse brains were rapidly excised, and the habenula was dissected and immediately submerged in RNAlater™ Solution (#AM7021, Invitrogen, Thermo Fisher Scientific, Carlsbad, CA, USA) to stabilise RNA. Samples were stored at 4 °C until processing. Total RNA was extracted using the miRNeasy Micro

Kit (#217084, QIAGEN Germantown, MD, USA) following the manufacturer's instructions. Complementary DNA (cDNA) was synthesised using the iScript cDNA Synthesis Kit (#1708891, Bio-Rad Laboratories, Hercules, CA, USA). Gene expression was quantified by real-time PCR (CFX Duet real-time PCR system, Bio-Rad Laboratories) using an iQ SYBR® Green Supermix (#1708882, Bio-Rad Laboratories). Each sample was analysed in triplicate technical replicates. Amplified specificity was confirmed by agarose gel electrophoresis. Relative gene expression was calculated using the comparative ΔΔCt method, and normalised to *Gapdh*. Fold changes in cytokine, *Ddc* and *Hcrtr2* expression were determined by comparing LH- or CRS-model mice with their respective controls. Primer sequences are provided in Supplementary Table 2.

**Human habenular expression analysis**

Archived post-mortem human habenular tissue was obtained from the Douglas Bell Canada Brain Bank (DBCBB; Douglas Mental Health University Institute, Verdun, Quebec, Canada). The cohort, tissue processing and habenula dissection procedures have been described previously[1]. For the present study, residual archived samples from the same previously characterised cohort were used for additional expression analysis of *DDC* and *HCRTR2* in the human habenula. The cohort comprised 12 suicide subjects with major depressive disorder (MDD) and 11 psychiatrically healthy control subjects. All subjects were male and of Caucasian ancestry, and the groups were matched for age, tissue pH and post-mortem interval.

Brains were hemisected, meninges removed, and sectioned coronally into 18–20 slabs before freezing. Habenular tissue was dissected from slabs 11–12 using a round-tipped burin. Total RNA (500 ng) was reverse-transcribed with random primers using iScript Reverse Transcription Supermix (Bio-Rad Laboratories). RNA purity and integrity were verified with a NanoDro p ND-

1000 spectrophotometer (Thermo Fisher Scientific). Stability of 28S rDNA and Gapdh reference gene expression was confirmed across groups. Quantitative real-time PCR was performed using iQ SYBR® Green Supermix (Bio-Rad Laboratories) on a CFX Duet system (Bio-Rad Laboratories). Reactions (20 µL) contained 2 µL cDNA, 10 µL SYBR® Green Supermix, and 0.5 µM of each primer. Thermal cycling conditions were: 95 °C for 3 min, followed by 40 cycles of 95 °C for 15 s, 60 °C for 1 min, and 60 °C for 30 s. Specificity was confirmed by melting curve analysis. All reactions were performed in triplicate. Relative gene expression was calculated using the ΔΔCt method with GAPDH as the internal control. Primer information is provided in Supplementary Table 3.

**DNA methylation analysis**

**Genomic DNA extraction and bisulfite conversion**

The LHb was microdissected from mice subjected to CRS or LH protocols. Genomic DNA was extracted using the Wizard® Genomic DNA Purification Kit (Promega, Madison, WI, USA) according to the manufacturer's instructions. DNA concentration and purity were assessed using a NanoDrop ND-1000 spectrophotometer (Thermo Fisher Scientific, Waltham, MA, USA). For bisulfite conversion, 2 µg genomic DNA was diluted in 20 µL ultrapure water (WELGENE, Gyeongsan, Republic of Korea). Bisulfite conversion was performed using the EpiJET Bisulfite Conversion Kit (Thermo Fisher Scientific) according to the manufacturer's protocol. Converted DNA was eluted in 50 µL elution buffer and stored at −80 °C until use.

**Methylation-specific PCR (MSP) and quantitative MSP (qMSP)**

MSP was performed using primers specific for methylated and unmethylated CpG sites within gene promoter regions, designed using MethPrimer. Primer sequences, product sizes and annealing temperatures are listed in Supplementary Table 4. PCR products generated from bisulfite-converted DNA were separated on 1.2% agarose gels and visualised using a FireReader V10 Gel Documentation System (Thistle Scientific Ltd, Glasgow, UK).

qMSP for *Ddc* was performed using a QuantStudio™ 3 Real-Time PCR System (Thermo Fisher Scientific, Waltham, MA, USA). Each 20-μL reaction contained 10 μL 2× Maxima SYBR Green/ROX qPCR Master Mix (Thermo Fisher Scientific), 250 nM of each primer, and 30 ng bisulfite-converted DNA. The thermal cycling conditions were 95 °C for 2 min, followed by 45 cycles of 95 °C for 15 s and 60 °C for 1 min. All reactions were performed in triplicate. Relative methylation levels were calculated using the ΔCt method and normalised to *Gapdh* or *Guca2* as internal reference genes. Probe information is provided in Supplementary Table 4.

**Pharmacological activation of OX2R**

To test whether activation of OX2R in the LHb promotes stress resilience, adult male $Ddc^{Cre}$ mice were stereotaxically implanted with 26-gauge dual guide cannulae (Plastics One, Roanoke, VA, USA) targeting the LHb bilaterally. After 1 week of recovery, mice underwent an LH protocol consisting of 100 inescapable footshocks per day for 3 consecutive days. On day 4, orexin-A (1 μL, 333 pmol $\mu L^{-1}$ in sterile saline containing 0.1% DMSO) or vehicle was unilaterally infused into the LHb through a 33-gauge internal injector inserted through the pre-implanted guide cannulae. Infusions were delivered at 100 nL $min^{-1}$ using an UltraMicroPump III, and the injector was left in place for 20 min after infusion to allow diffusion and minimise backflow. Mice were then returned to their home cages and underwent behavioural testing 4 days later in the OFT, followed by the TST on the next day.

To assess whether non-invasive restoration of orexin signalling could rescue stress-induced deficits, adult male *Ddc*$^{Cre}$ mice subjected to the LH protocol received intranasal orexin-A administration on day 4, 1 day after completion of the LH procedure. Orexin-A was prepared at 6 μg μL$^{-1}$ in sterile saline containing 0.1% DMSO. Mice received alternating intranasal administrations of 1 μL per nostril, with an interval of approximately 30 s between nostrils, until a total volume of 10 μL had been delivered (five administrations per nostril). During administration, mice were gently restrained in a supine position to facilitate absorption through the nasal mucosa. Behavioural testing was performed 4 days later in the OFT, followed by the TST on the next day and the FST on the following day.

To determine whether AMP-activated protein kinase (AMPK) contributes to orexin-A-dependent rescue, mice subjected to the LH protocol received Compound C (CC) pretreatment before intranasal orexin-A administration. CC (10 mg kg$^{-1}$; #171260, Calbiochem, Merck KGaA, Darmstadt, Germany) was first dissolved in DMSO, then mixed with Tween-80 and sterile saline, and sonicated to obtain a uniform injectable suspension. Vehicle-treated mice received the corresponding solvent mixture without CC. CC or vehicle was administered by intraperitoneal injection 1 h before intranasal orexin-A treatment. Behavioural testing was performed 4 days later in the OFT, followed by the TST on the next day and the FST on the following day.

For molecular analyses, mice were killed after completion of behavioural testing, and the habenula was microdissected by tissue punching for qPCR. For FISH and histological analyses, mice were transcardially perfused, and brains were collected and processed as described above. All behavioural scoring and data analysis were performed by experimenters blinded to treatment allocation.

**Statistical analyses**

All data are presented as mean ± s.e.m. Statistical analyses were performed using GraphPad Prism 10 (GraphPad Software, Boston, MA, USA), unless otherwise stated. Normality was assessed using the Shapiro–Wilk test. For comparisons between two groups, two-tailed unpaired or paired Student's *t*-tests were used as appropriate. For comparisons involving more than two groups or repeated measurements, one-way ANOVA, repeated-measures one-way ANOVA, or two-way ANOVA was used as appropriate, followed by the *post hoc* multiple-comparisons test indicated in the corresponding figure legend or Supplementary Table 5. Statistical significance was defined as $P < 0.05$.

Sample sizes are indicated in the corresponding figure legends and in Supplementary Table 5. In Supplementary Table 5, sample-size information is reported in the **n** column according to experiment type. For behavioural, pharmacological, qPCR, methylation and other animal-based experiments, the value in the **n** column indicates the number of biologically independent mice. For electrophysiological, FISH and immunohistochemical analyses, the **n** column reports the number of independent animals together with the number of quantified cells, neurons, sections or imaging fields in parentheses, as specified in the corresponding figure legend or table entry. For qPCR, technical replicates were averaged to obtain a single value for each biological sample before statistical analysis. Mice were randomly assigned to experimental groups, and data acquisition and analysis were performed blind to group allocation whenever feasible. Data were excluded only based on pre-established technical criteria described above, including mistargeted viral expression or fibre placement, unstable recordings, or predefined electrophysiological quality-control thresholds. All key experiments were independently replicated at least twice with consistent results. Sample sizes were estimated on the basis of prior literature, preliminary effect sizes and feasibility; no formal statistical methods were used to predetermine sample size.

### Figure legends

**Extended Data Fig. 1 Anatomical and functional validation of LHA inputs to LHb D-neurons.**

**a**, Representative images of starter cells in the LHb of *Ddc*$^{\mathrm{Cre}}$ mice used for monosynaptic retrograde tracing with Cre-dependent TVA/RG helper AAVs and RVΔG. RVΔG-labelled cells are shown in green and TVA-expressing starter cells in red; bottom, enlarged views of the boxed regions. **b**,**c**, Atlas-based mapping of RVΔG-labelled presynaptic neurons across representative coronal sections, showing distributed inputs to LHb D-neurons from forebrain and hypothalamic regions, including the MS, LS, LHA and LPO. **d**, Representative images showing RVΔG-labelled presynaptic neurons in the LHA together with FISH detection of *Hcrt* and *Vglut2*. **e**–**g**, Higher-magnification images of RVΔG signal (**e**), *Hcrt* (**f**) and *Vglut2* (**g**). **h**,**i**, Merged images showing co-localisation of RVΔG-labelled neurons with *Hcrt* (**h**) and *Vglut2* (**i**). **j**, Triple-merged image demonstrating that RVΔG-labelled input neurons co-express *Hcrt* and *Vglut2*. These findings indicate that presynaptic inputs from the LHA to LHb D-neurons include orexin/glutamatergic neurons. **k**,**l**, Representative voltage-clamp traces from LHb neurons showing light-evoked inward currents in response to optogenetic stimulation of LHA orexin terminals (**k**), and expanded representative trace showing inward currents evoked by repeated blue-light stimulation (**l**). Blue dashed lines indicate light pulses. These data support direct functional connectivity between LHA orexin terminals and LHb neurons. AAV, adeno-associated virus; RVΔG, glycoprotein-deleted rabies virus; TVA, avian tumour virus receptor A; RG, rabies glycoprotein; LHb, lateral habenula; LHA, lateral hypothalamic area; MS, medial septum; LS, lateral septum; LPO, lateral preoptic area; FISH, fluorescence *in situ* hybridization; *Hcrt*, hypocretin; *Vglut2*, vesicular glutamate transporter 2. Scale bars, **a**: 50 µm, **a'**–**a''''**: 10 µm, **b**,**c**: 500 µm, **d**: 50 µm, **e**–**j**: 20 µm.

**Extended Data Fig. 2 Chemogenetic activation of LHb-projecting orexin neurons does not alter open-field locomotion.**

**a**, Representative images of dopamine sensor GRAB-rDA3m expression in the NAc. **b**, Schematic showing the placement of the fibre photometry optic fibre in the NAc used in the experiment reported in Fig. 2**g**,**h**. **c**–**h**, OFT measures in *Orexin*$^{\mathrm{Cre}}$ mice after projection-defined chemogenetic activation of LHb-projecting orexin neurons. Quantification of total distance moved across groups (**c**), paired comparisons of total distance moved before and after CNO administration in mCherry control mice (**d**) and hM3Dq mice (**e**), quantification of mean velocity across groups (**f**), and paired comparisons of mean velocity before and after CNO administration in mCherry control mice (**g**) and hM3Dq mice (**h**). Chemogenetic activation did not significantly alter locomotor activity, indicating that the behavioural effects observed in Fig. 2 were not attributable to a general increase in movement. NAc, nucleus accumbens; CNO, clozapine N-oxide; OFT, open field test. Error bars indicate mean ± s.e.m. NS, not significant. Scale bars, **a**: 500 μm. Exact *n* and *P* values are provided in Supplementary Table 5.

**Extended Data Fig. 3 Optogenetic activation of LHA orexin terminals in the LHb rapidly and reversibly promotes coping and positive valence.**

**a**, Experimental schematic for optogenetic stimulation of LHA orexin terminals in the LHb of *Orexin*$^{\mathrm{Cre}}$ mice during behavioural testing. **b**, Representative images showing EYFP-labelled orexin projections in the LHb, optical fibre placement above the LHb, and overlap between orexin immunoreactivity and EYFP-labelled terminals. **c**,**d**, Temporal profile of behaviour during the TST under an OFF–ON–OFF stimulation design (**c**), and quantification across epochs showing increased struggling during photostimulation in ChR2 mice but not in EYFP controls (**d**). **e**,**f**,

Representative heat maps from the RTPP assay showing preference for the photostimulation-paired chamber in ChR2 mice (**e**), and quantification of time spent in the stimulation-paired chamber (**f**). **g**,**h**, OFT measures showing total distance moved across time under alternating stimulation epochs (**g**), and quantification of total distance moved across groups (**h**). **i**,**j**, RTPP measures showing total distance moved (**i**) and mean velocity (**j**) across groups. Photostimulation rapidly increased active coping and induced real-time place preference without significantly altering OFT or RTPP locomotor measures. LHA, lateral hypothalamic area; LHb, lateral habenula; TST, tail suspension test; RTPP, real-time place preference; OFT, open field test; ChR2, channelrhodopsin-2. Error bars indicate mean ± s.e.m. NS, not significant; $^{**}P < 0.01$, $^{****}P < 0.0001$. Scale bars, **b**: 100 μm. Exact *n* and *P* values are provided in Supplementary Table 5.

**Extended Data Fig. 4 Orexin-B does not reproduce the OX2R-sensitive inhibitory profile of orexin-A in LHb neurons.**

**a**–**d**, EPSC frequency (**a**), normalised EPSC frequency (**b**), EPSC amplitude (**c**) and normalised EPSC amplitude (**d**) in LHb neurons recorded in the presence of bicuculline and picrotoxin, showing that orexin-B did not significantly alter excitatory synaptic transmission and did not produce a TCS-OX2-29-sensitive effect. **e**,**f**, Representative voltage-clamp trace showing an orexin-B-induced inward current (**e**), and quantification of inward current amplitude relative to baseline (**f**). **g**,**h**, Representative current-clamp traces showing that orexin-B increased spike output in response to depolarising current injection (**g**), and quantification of the number of action potentials across injected current steps under control and orexin-B conditions (**h**). **i**,**j**, Representative current-clamp traces showing that the orexin-B-induced increase in firing was further enhanced in the presence of $Ba^{2+}$ (**i**), and quantification of firing rate across baseline, orexin-B and orexin-B+$Ba^{2+}$ conditions (**j**). These data indicate that, unlike orexin-A, orexin-B did not measurably suppress excitatory synaptic strength in LHb neurons and instead increased neuronal firing while exerting a distinct membrane effect. LHb, lateral habenula; EPSC, excitatory postsynaptic current; OX2R, orexin receptor 2. Error bars indicate mean ± s.e.m. NS, not significant; $^{**}P < 0.01$, $^{***}P < 0.001$, $^{****}P < 0.0001$. Exact *n* and *P* values are provided in Supplementary Table 5.

**Extended Data Fig. 5 *In vivo* recording of LHb-projecting LHA orexin neurons**

**a**, Characteristics of light-evoked responses in $Orexin^{Cre}$ mice. Peristimulus time histograms (10-ms bins) constructed during the presentation of 10 trains of 10 blue-light pulses (each pulse 5-ms width at 30 Hz; inter-train interval of 60 sec). Spike probability and latency in response to 100 pulses were calculated from all LHA neurons recorded in each group. The boxed histograms show firing patterns of two representative orexin neurons. **b**, Correlations between spontaneous and light-evoked waveforms of orexin neurons. **c**, Average firing rates of orexin neurons (5-min bins) before, during and after tail suspension. LHA, lateral hypothalamic area; LHb, lateral habenula; ChR2, channelrhodopsin-2. Error bars indicate mean ± s.e.m. NS, not significant; $^{****}P < 0.0001$. Exact *n* and *P* values are provided in Supplementary Table 5.

**Extended Data Fig. 6 Chronic restraint stress similarly disrupts the orexin-responsive LHb D-neuron module.**

**a**–**e**, Behavioural and methylation analyses in the LH model, showing total distance moved (**a**), mean velocity (**b**), centre duration in the OFT (**c**), immobility time in the FST (**d**), and methylation-specific PCR images for *Ddc* and *Hcrtr2* (**e**). **f**, Experimental schematic of the CRS protocol. **g**–**j**, Behavioural analyses in the CRS model, showing total distance moved (**g**), mean velocity (**h**), centre duration in the OFT (**i**), and immobility time in the FST (**j**). **k**–**o**, Representative *Fos* images and quantification showing increased activity in LHA orexin neurons (**k**), LHb D-neurons (**l**), increased activity in RMTg GABAergic neurons (**m**), and reduced activity in VTA dopaminergic neurons (**n**) and NAc GABAergic neurons (**o**) after CRS. **p**,**q**, Representative FISH images and quantification showing reduced *Tet2* signal within *Ddc*$^{+}$ neurons (**p**) and reduced *Hcrtr2* signal within *Ddc*$^{+}$ neurons (**q**) after CRS. **r**–**v**, qPCR analysis of LHb transcripts after CRS showing changes in cytokine and D-neuron-related gene expression, including *Il1b* (**r**), *Tnf* (**s**), *Il6* (**t**), *Ddc* (**u**) and *Hcrtr2* (**v**). **w**–**y**, Methylation-specific PCR images for *Ddc* and *Hcrtr2* after CRS (**w**), and quantification of *Ddc* (**x**) and *Hcrtr2* (**y**) promoter methylation in control and CRS mice. **z**, Working model summarising the circuit-wide shift under chronic stress, with increased activity in the LHA, LHb and RMTg, and reduced activity in the VTA and NAc. LHA, lateral hypothalamic area; LHb, lateral habenula; RMTg, rostromedial tegmental nucleus; VTA, ventral tegmental area; NAc, nucleus accumbens; LH, learned helplessness; CRS, chronic restraint stress; OFT, open field test; FST, forced swim test; qPCR, quantitative polymerase chain reaction; FISH, fluorescence *in situ* hybridization; Il1b, interleukin-1β; Tnf, tumour necrosis factor; Il6, interleukin-6; Ddc, dopa decarboxylase; Hcrtr2, hypocretin receptor 2; Tet2, ten-eleven translocation 2; GLU, glutamate; OX-B, orexin-B; DA, dopamine. Error bars indicate mean ± s.e.m. NS, not significant; *$P < 0.05$, **$P < 0.01$, ***$P < 0.001$, ****$P < 0.0001$. Exact *n* and *P* values are provided in Supplementary Table 5.

**Extended Data Fig. 7 Human post-mortem habenular *HCRTR2* expression is relatively preserved in MDD.**

**a**–**d**, Human post-mortem habenula analysis of *HCRTR2*, including comparison of *HCRTR2* expression between control and MDD cases who died by suicide (**a**), distribution analysis of transformed *HCRTR2* expression values (**b**), group comparison of transformed *HCRTR2* expression by Welch's *t*-test (**c**), and heat map of human habenular *HCRTR2* expression (**d**). MDD, major depressive disorder; Hcrtr2, hypocretin receptor 2. Error bars indicate mean ± s.e.m. NS, not significant. Exact *n* and *P* values are provided in Supplementary Table 5.

**Extended Data Fig. 8 Intra-LHb orexin-A rescue and behavioural effects of D-neuron-specific *Hcrtr2* knockdown.**

**a**, Experimental schematic showing the LH protocol, followed by bilateral intra-LHb infusion of orexin-A or vehicle through pre-implanted cannulae one day after completion of LH; behavioural testing was performed 4 days after infusion. **b**, Quantification of immobility time in the TST. **c**–**g**, qPCR analysis of LHb transcripts showing changes in *Il1b* (**c**), *Tnf* (**d**), *Il6* (**e**), *Ddc* (**f**) and *Hcrtr2* (**g**) after LH and intra-LHb orexin-A treatment. **h**–**j**, OFT measures showing total distance moved (**h**), mean velocity (**i**) and centre duration (**j**). Intra-LHb orexin-A reduced immobility in the TST and altered selected LH-induced molecular readouts without significantly affecting OFT measures. **k**–**n**, Behavioural analyses following Cre-dependent knockdown of *Hcrtr2* in LHb D-neurons, showing total distance moved (**k**), mean velocity (**l**), centre duration in the OFT (**m**), and immobility time in the FST (**n**). *Hcrtr2* knockdown did not significantly alter OFT measures but increased immobility in the FST, consistent with a contribution of endogenous OX2R signalling to active coping behaviour. LHb, lateral habenula; LH, learned helplessness; TST, tail suspension test; OFT, open field test; qPCR, quantitative polymerase chain reaction; *Il1b*, interleukin-1β; *Tnf*, tumour necrosis factor; *Il6*, interleukin-6; *Ddc*, dopa decarboxylase; *Hcrtr2*, hypocretin receptor

2. Error bars indicate mean ± s.e.m. NS, not significant; *$P < 0.05$, **$P < 0.01$, ***$P < 0.001$, ****$P < 0.0001$. Exact *n* and *P* values are provided in Supplementary Table 5.

**Extended Data Fig. 9 Compound C attenuates orexin-A-dependent rescue after LH.**

**a**, Experimental schematic showing the LH protocol, followed by intraperitoneal injection of CC 1 h before intranasal orexin-A administration; behavioural testing was performed 4 days later. **b**–**d**, OFT measures showing total distance moved (**b**), mean velocity (**c**) and centre duration (**d**). **e**,**f**, Quantification of immobility time in the TST (**e**) and FST (**f**). **g**–**k**, qPCR analysis of LHb transcripts showing changes in *Il1b* (**g**), *Tnf* (**h**), *Il6* (**i**), *Ddc* (**j**) and *Hcrtr2* (**k**) after CC pretreatment and intranasal orexin-A administration. **l**, Representative images showing p-NF-κB (green) in mCherry-labelled LHb D-neurons (red) across conditions. **m**,**n**, Representative images of *Tet2* expression in LHb cells (**m**), and quantification of the proportion of $Tet2^{+}$ cells in the LHb (**n**). CC pretreatment altered OFT performance, attenuated the behavioural effects of orexin-A, and reversed orexin-A-dependent molecular rescue in LHb D-neurons. LH, learned helplessness; CNO, clozapine N-oxide; OFT, open field test; TST, tail suspension test; FST, forced swim test; qPCR, quantitative polymerase chain reaction; *Il1b*, interleukin-1β; *Tnf*, tumour necrosis factor; *Il6*, interleukin-6; *Ddc*, dopa decarboxylase; *Hcrtr2*, hypocretin receptor 2; NF-κB, nuclear factor κB; *Tet2*, ten-eleven translocation 2. Error bars indicate mean ± s.e.m. NS, not significant; *$P < 0.05$, **$P < 0.01$, ****$P < 0.0001$. Scale bars, **i**,**m**: 10 μm. Exact *n* and *P* values are provided in Supplementary Table 5.

**Extended Data Fig. 10 Proposed model of a self-limiting orexin–habenula resilience circuit.**

Schematic summary integrating the present findings with prior literature. LHA orexin/glutamatergic neurons engage OX2R-enriched D-neurons in the LHb, which in turn suppress RMTg GABAergic neurons and promote VTA dopaminergic output and NAc recruitment. Feedback through the broader LHA→LHb→RMTg→VTA→NAc→LHA network is proposed to constrain this coping-promoting state after stress, consistent with the biphasic activity dynamics of LHb-projecting orexin neurons observed *in vivo*. At the cellular level, orexin-A is proposed to engage two functionally distinguishable components in LHb D-neurons: an acute OX2R-sensitive inhibitory component compatible with a $Ba^{2+}$-sensitive inwardly rectifying $K^+$ conductance, and a slower protective component in which OX2R-associated Gαq–PLCβ–$Ca^{2+}$ signalling activates CaMKKβ–AMPK, restrains NF-κB signalling, preserves *Tet2*, and supports demethylation-linked maintenance of the *Ddc* locus. Consistent with prior receptor literature indicating that OX2R can couple to both PTX-sensitive Gαi/o and PTX-insensitive Gαq pathways, the present LHb data support a model in which both signalling modes may operate in this circuit, with the inhibitory component linked to OX2R-sensitive inwardly rectifying $K^+$ conductance and the protective component linked to Gαq–$Ca^{2+}$–AMPK signalling. Under chronic stress, this resilience module is proposed to collapse in association with inflammatory activation, promoter methylation and erosion of D-neuron identity and orexin responsiveness. The ketamine annotation highlights a potential point of functional convergence at the level of LHb hyperactivity control, rather than a shared upstream receptor mechanism. Black arrows indicate established or proposed circuit flow; red annotations denote inhibitory/protective signalling; blue annotations denote excitatory/plasticity-related signalling. This figure is a working model and does not imply that all intermediate signalling steps were directly tested in the present study.

**Supplementary Table 1. RNAscope FISH probes used in this work.**

| Gene name | Accession # | Target region | Cat. # | Manufacturer |
|---|---|---|---|---|
| *Ddc* | NM_001190448.1 | 297~1726 | 464871 | ACD (Advanced cell Diagnostics) |
| | | 297~1726 | 464871-C3 | |
| *Fos* | NM_010234.2 | 407~1427 | 316921 | |
| | | 407~1427 | 316921-C3 | |
| *Hcrt* | NM_010410.2 | 2~577 | 490461-C2 | |
| *Hcrtr2* | NM_198962.3 | 35~1085 | 460881 | |
| *Slc17a6* (*Vglut2*) | NM_080853.3 | 1986-2998 | 319171-C3 | |
| *Gad1* | NM_0080774 | 62~3113 | 400951-C3 | |
| *Tet2* | NM_001040400.2 | 401~1373 | 511591-C2 | |
| *Th* | NM_009377.1 | 483~1603 | 317621-C3 | |

**Supplementary Table 2. Primer sequences used for qPCR.**

| Gene name | Primer sequence (5' to 3') | | Location | Product size(bp) | Annealing (℃) | Reference |
|---|---|---|---|---|---|---|
| *Il1β* | Forward | CTGTGTCTTTCCCGTGGACC | 317 ~ 516 | 200 | 60 | NM_008361.4 |
| | Reverse | CAGCTCATATGGGTCCGACA | | | | |
| *Tnf* | Forward | CTGTAGCCCACGTCGTAGCA | 439 ~ 636 | 198 | 60 | NM_013693.3 |
| | Reverse | TGTGGGTGAGGAGCACGTAG | | | | |
| *Il6* | Forward | GAGGATACCACTCCCAACAGACC | 187 ~ 327 | 141 | 60 | NM_031168.2 |
| | Reverse | AAGTGCATCATCGTTGTTCATACA | | | | |
| *Ddc* | Forward | GGCTTACATCCGAAAGCACG | 1197 ~ 1314 | 118 | 60 | NM_016672.5 |
| | Reverse | CTTTAGCCGGAAGCAGACCA | | | | |
| *Hcrtr2* | Forward | GCAACTGGTCATCTGCTTCA | 157 ~ 329 | 173 | 60 | NM_198962.4 |
| | Reverse | GATGAGAGCCACAACGAACA | | | | |
| *Gapdh* | Forward | ACCCAGAAGACTGTGGATGG | 616 ~ 786 | 171 | 60 | NM_008084.4 |
| | Reverse | CACATTGGGGGTAGGAACAC | | | | |

**Supplementary Table 3. Primer information for human habenular *DDC* and *HCRTR2* expression analysis.**

| Gene name | Primer sequence (5' to 3') | | Location | Product size(bp) | Annealing (°C) | Reference |
|---|---|---|---|---|---|---|
| *DDC* | Forward | TGTGGAAGTCATTCTGGGGC | 1352 ~ 1507 | 156 | 60 | NM_001082971.2 |
| | Reverse | CGAGAACAGATGGCAAAGCG | | | | |
| *HCRTR2* | Forward | CCACCGACTATGACGACGAG | 165 ~ 289 | 125 | 60 | NM_001384272.1 |
| | Reverse | GTTCCCAATGAGAGCCACGA | | | | |
| *GAPDH* | Forward | ACCCACTCCTCCACCTTTGAC | 944 ~ 1053 | 110 | 60 | NM_002046.7 |
| | Reverse | TCCACCACCCTGTTGCTGTAG | | | | |

**Supplementary Table 4. Primers for MSP and qMSP.**

| Gene name | Primer sequence (5' to 3') | | Location | Product size(bp) | Annealing (℃) | Reference |
|---|---|---|---|---|---|---|
| *Ddc* | Me-F | GTTTTTGTCGGTTTTTGTTAGATTC | −2731 ~ −2601 | 131 | 60 | NC_000077.7 |
| | Me-R | TTATACCGAAATCTATTTTCAACGC | | | | |
| | Un-F | TTTTGTTGGTTTTTGTTAGATTTGT | −2729 ~ −2602 | 128 | 60 | |
| | Un-R | TATACCAAAATCTATTTTCAACACC | | | | |
| *Hcrtr2* | Me-F | TTGAAGTAGTAGTTCGAAGTTGTCG | −68 ~ +67 | 136 | 60 | NC_000075.7 |
| | Me-R | CATAAACACTAAAAATCCGCGAT | | | | |
| | Un-F | TGAAGTAGTAGTTTGAAGTTGTTGG | −67 ~ +69 | 137 | 60 | |
| | Un-R | CCCATAAACACTAAAAATCCACAAT | | | | |
| *ACTB* | Me-F | TTTGGGTAAGTTGTAGTTTAG | +2407 ~ +2524 | 118 | 60 | NC_000071.7 |
| | Me-R | TACCACAAAATTCCATACCTA | | | | |
| *Guca2* | Me-F | GGTGTTGTGGTTTAGAAGGTTATGG | −2555 ~ −2471 | 85 | 60 | ref[3] |
| | Me-R | ACCTTATCCTCAACTTCCAACATACC | | | | |

Me-F, methylated forward primer; Me-R, methylated reverse primer, Un-F, unmethylated forward primer; Un-F, unmethylated reverse primer
