## Supplementary figures and images for "A self-limiting orexin–habenula circuit for stress resilience"

### Extended Data figure 1.pdf

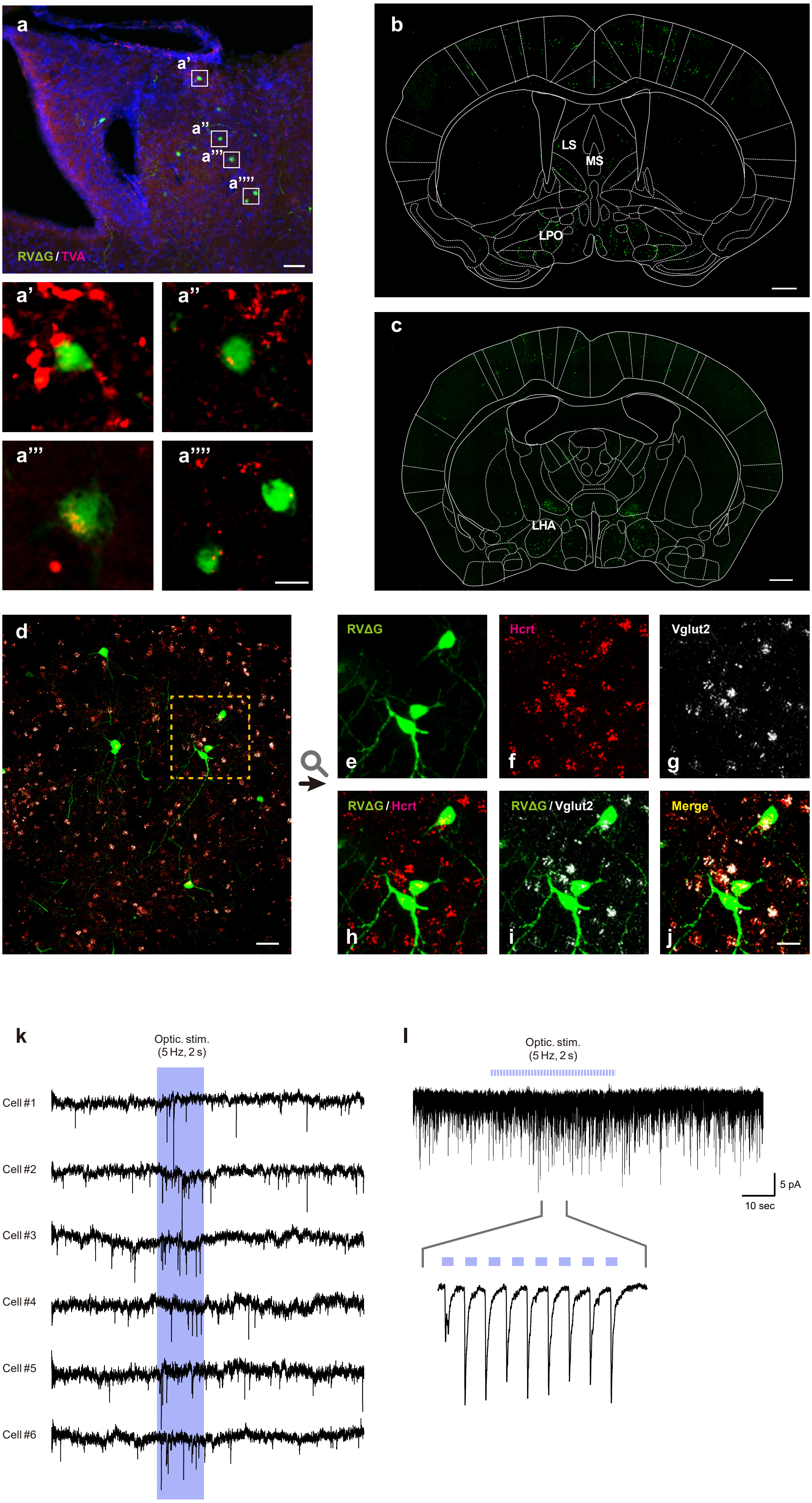

### Extended Data figure 2.pdf

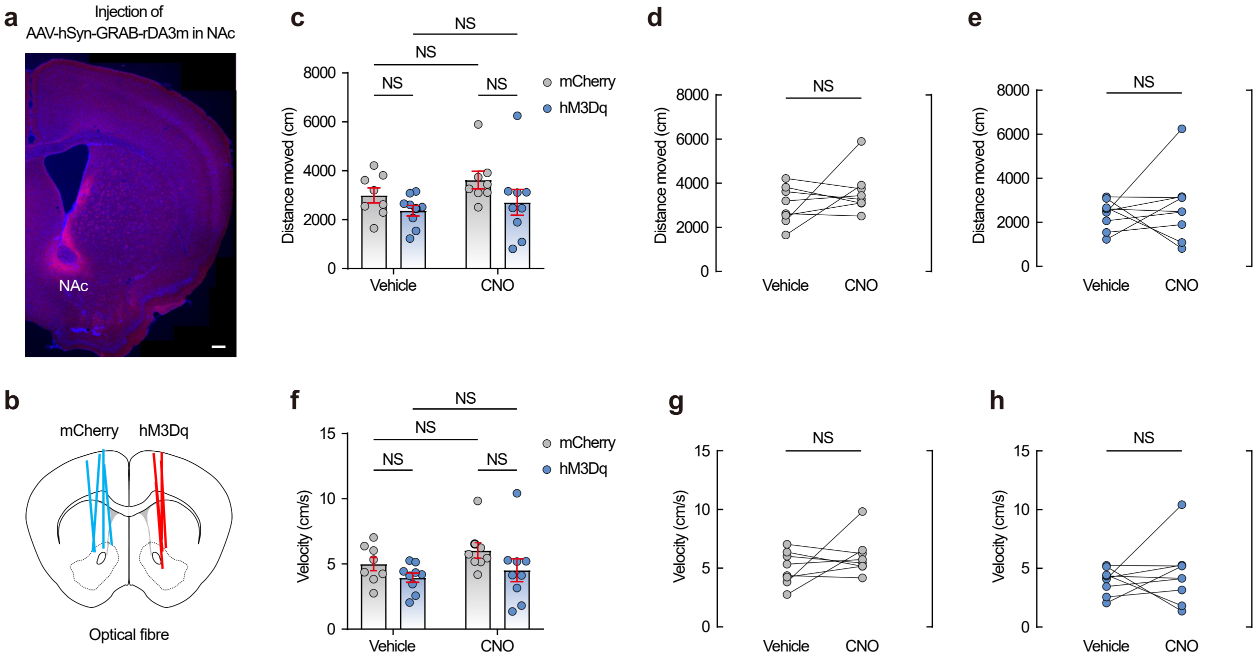

### Extended Data figure 3.pdf

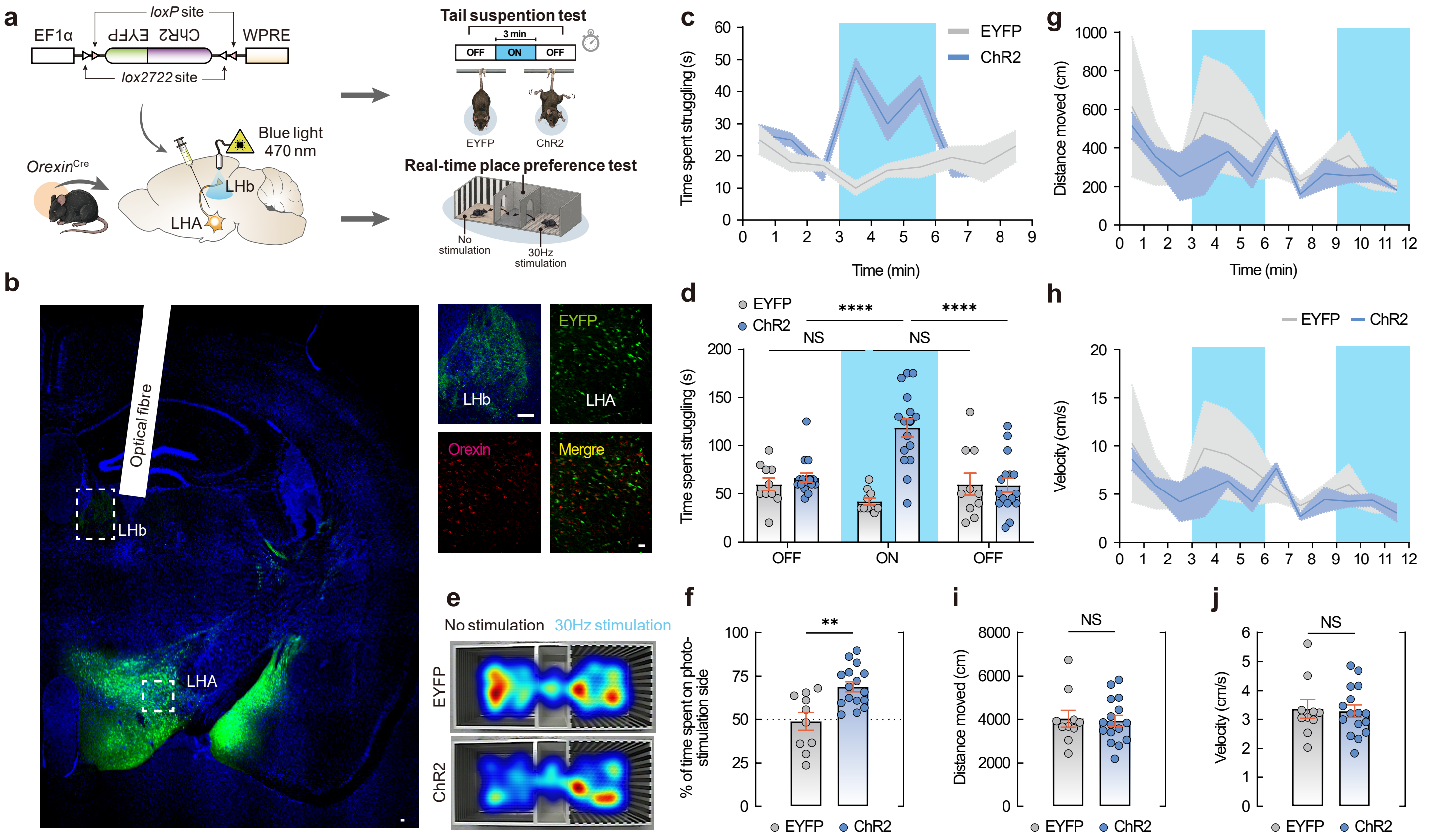

### Extended Data figure 4.pdf

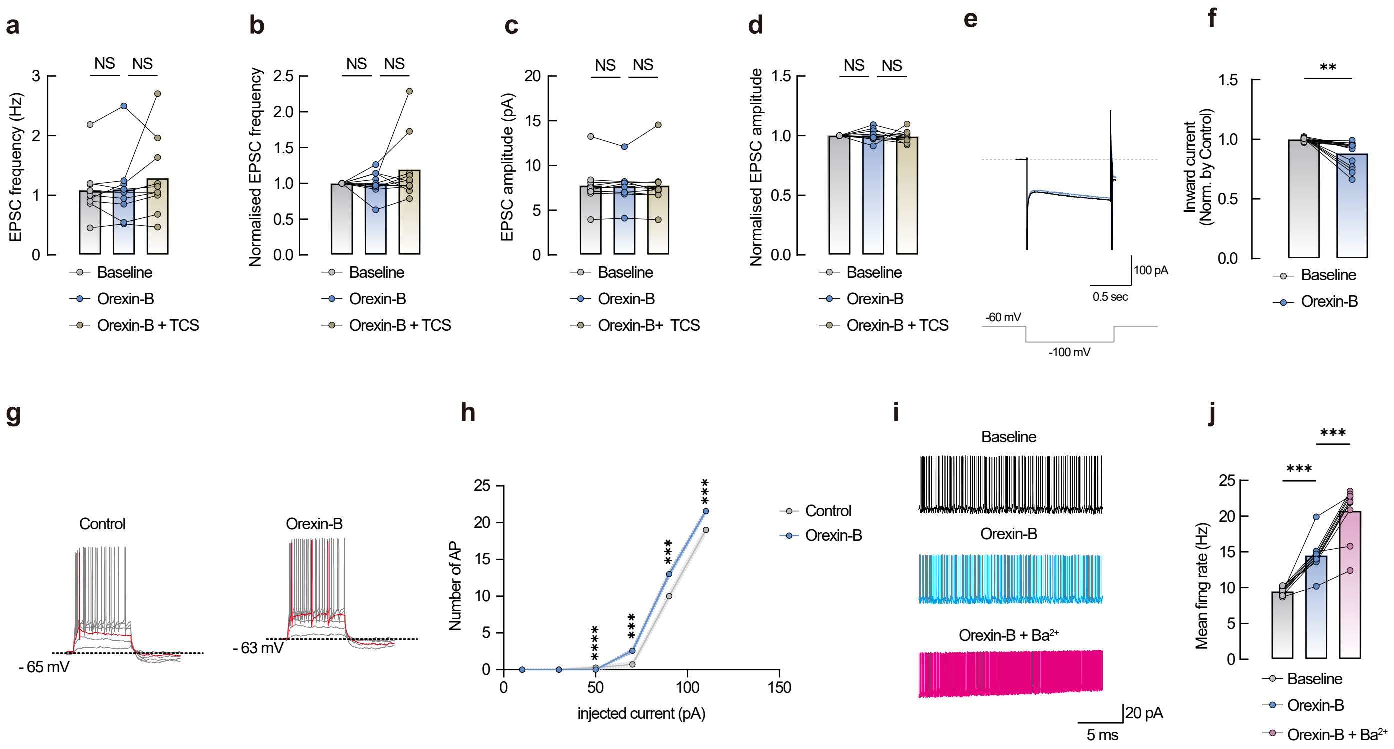

### Extended Data figure 5.pdf

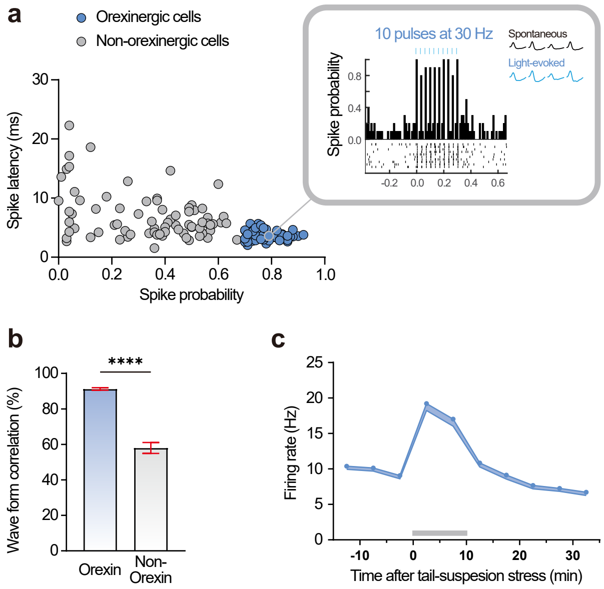

### Extended Data figure 6.pdf

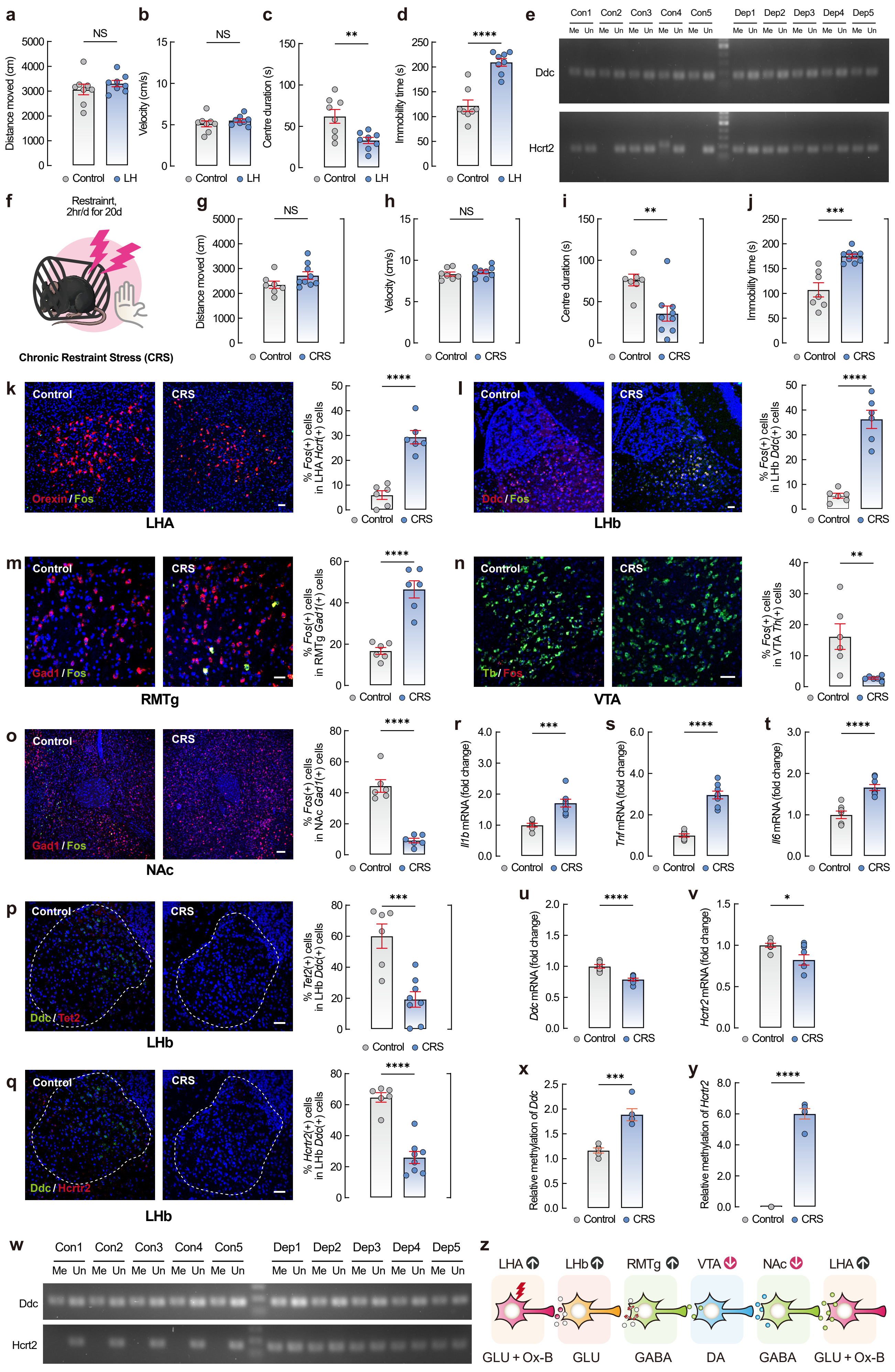

### Extended Data figure 7.pdf

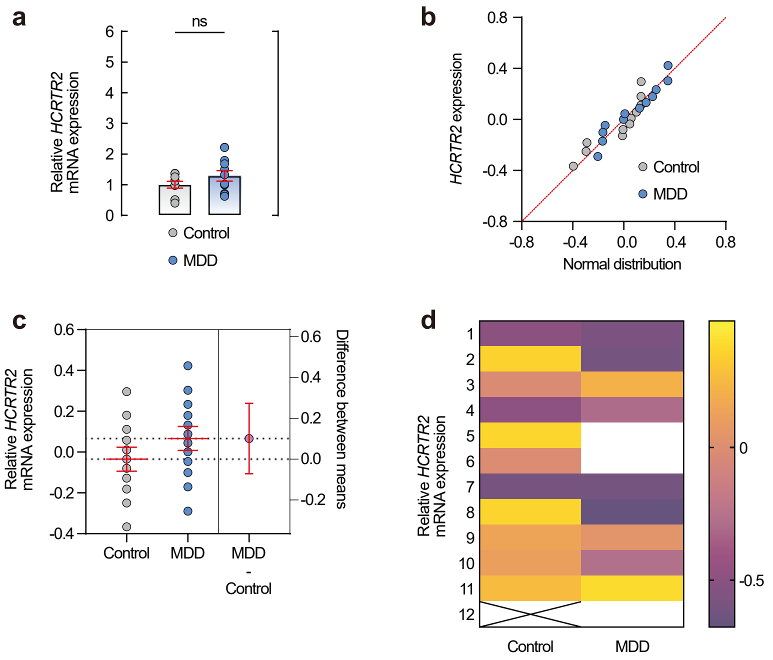

### Extended Data figure 8.pdf

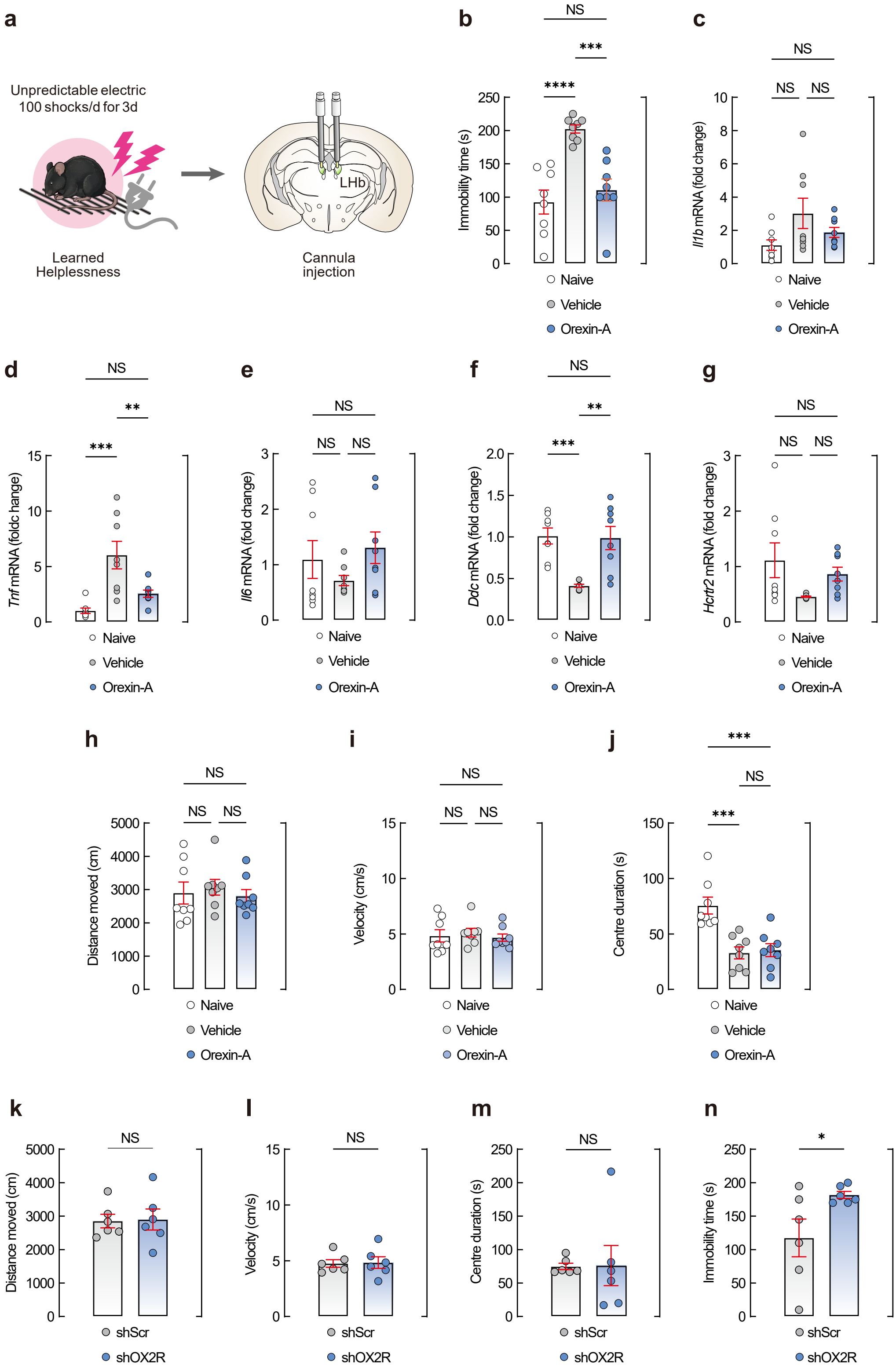

### Extended Data figure 9.pdf

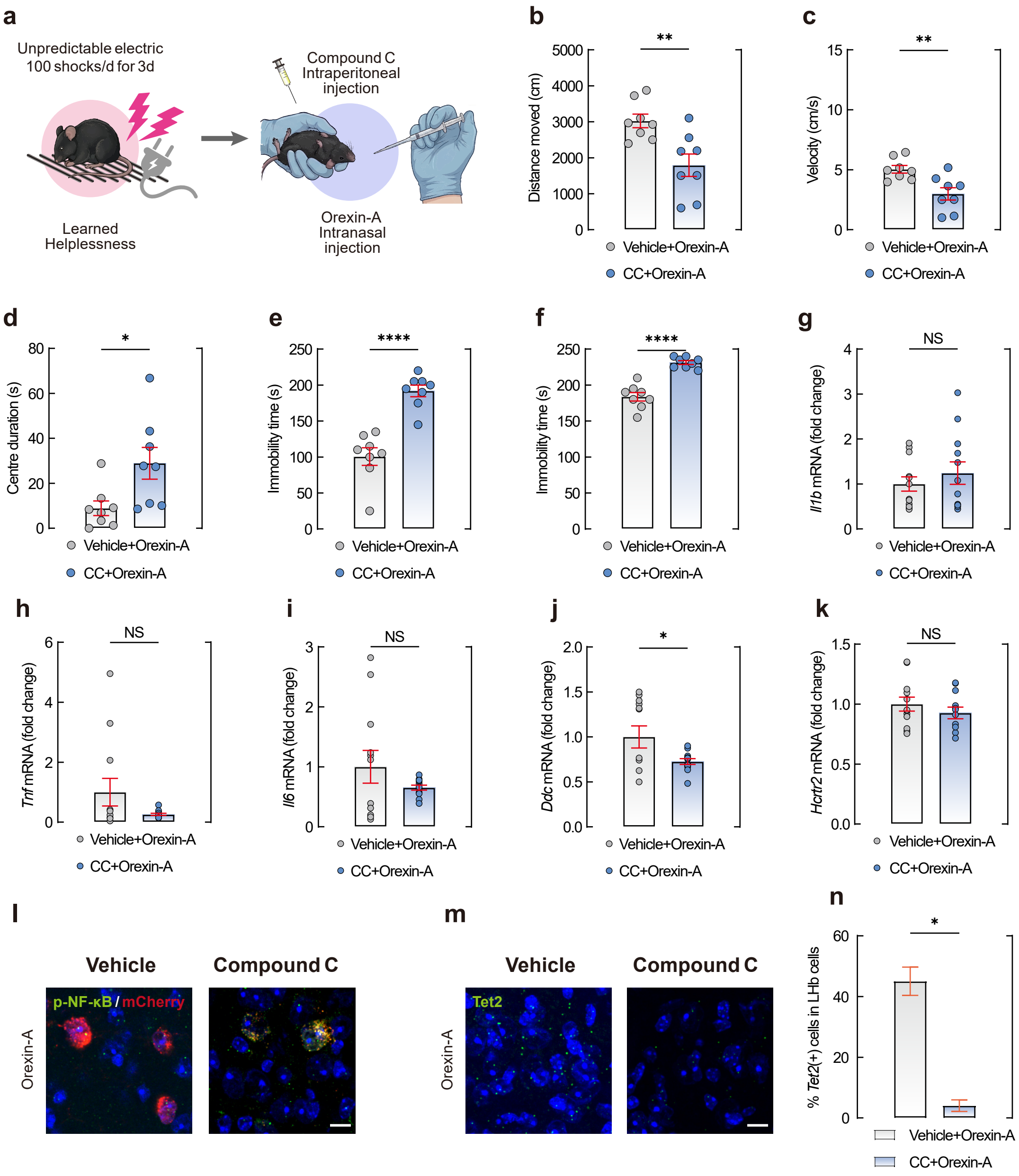

### Extended Data figure 10.pdf

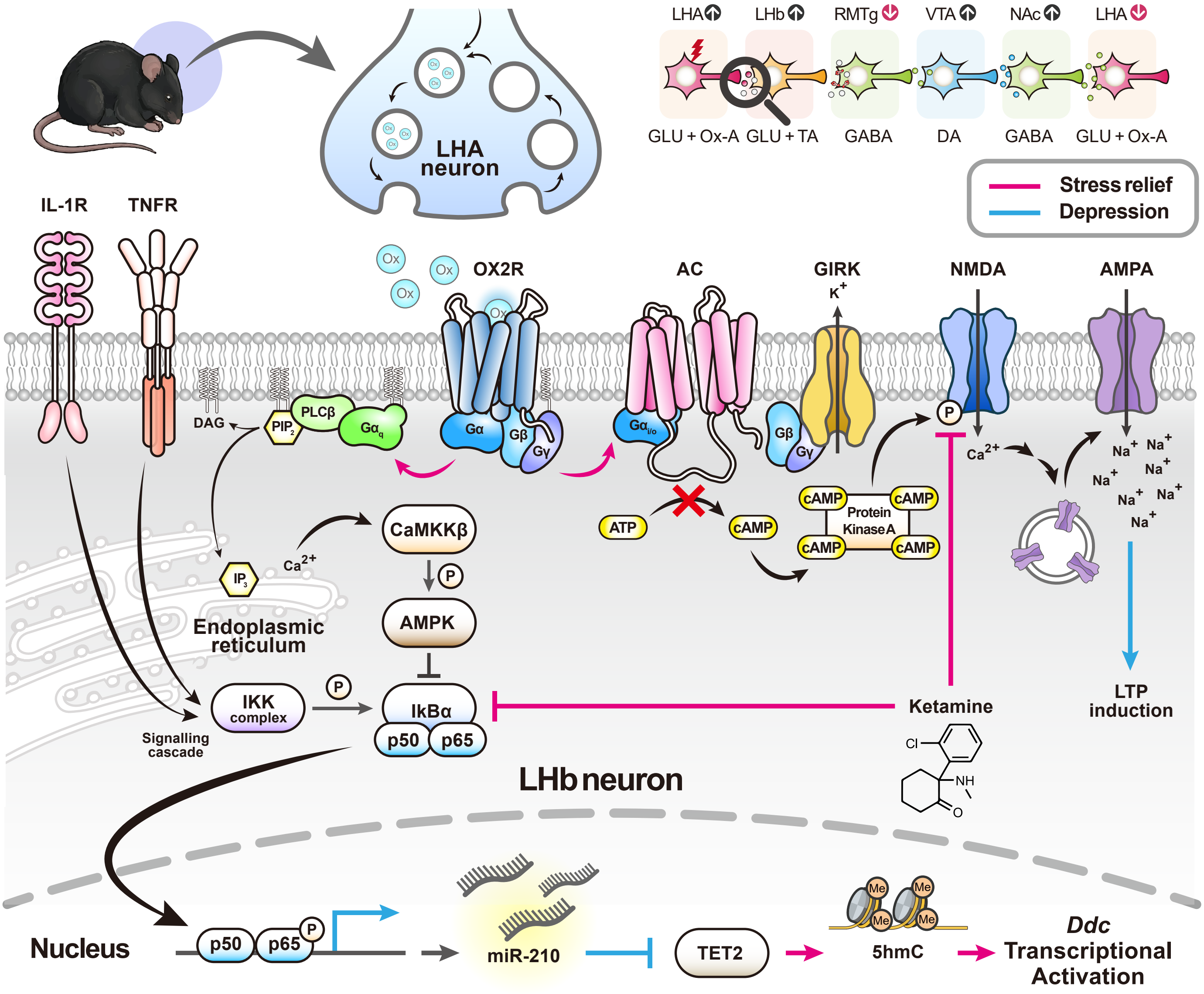
